# A non-canonical planar cell polarity pathway triggered by light

**DOI:** 10.1101/2021.04.21.440771

**Authors:** Michael Housset, Dominic Filion, Nelson Cortes, Hojatollah Vali, Craig Mandato, Christian Casanova, Michel Cayouette

## Abstract

The coordinated spatial arrangement of organelles within a tissue plane, generally referred to as Planar Cell Polarity (PCP), is critical for various processes ranging from organogenesis to sensory detection. Mechanistically, planar cell polarity is typically set-up by gradients of morphogens and their receptors, but whether non-molecular cues, akin to phototropism in plants, might play a part in setting up PCP in animals remains unknown. Here we report that the basal body of newborn photoreceptor cells in the mouse retina is centrally positioned, but then moves laterally around the first post-natal week, thereby generating cell-intrinsic asymmetry. During the second postnatal week, when eyes open, the basal bodies of cone photoreceptor cilia, but not rods, become coordinated across the tissue plane to face the center of the retina. Mechanistically, we show that light is required during a critical window in development, triggering a non-canonical cascade whereby cone transducin interacts with the G-protein signaling modulator protein 2 (GPSM2), which is in turn necessary for cone PCP. This work uncovers a non-canonical PCP pathway initiated by light.

## INTRODUCTION

In addition to apicobasal polarity, epithelial cells often display coordinated alignment of cell-intrinsic asymmetries across the tissue plane, an organization known as planar cell polarity (PCP). Striking examples of PCP are found in *Drosophila*, such as the uniform alignment of sensory bristles along the body wall or the precise alignment of ommatidia containing different photoreceptor cell types arranged in a stereotyped fashion across the eye^1–3^. In the latter, PCP is essential for tissue patterning, but may also optimize visual perception by facilitating synaptic convergence of identically oriented light-sensing photoreceptor cells to the brain, a wiring process known as neuronal superposition^4,5^. In the mammalian retina, rod and cone photoreceptors detect dim or bright light, respectively, with cones additionally sensitive to specific wavelengths. In both rods and cones, the photosensitive cilia, or outer segment (OS), develops from the connecting cilium, a microtubule-based structure anchored by a basal body to the apical surface of the inner segment (IS), which houses cellular organelles^6^. While the mechanisms and role of photoreceptor apicobasal polarity in vision have been extensively studied^7,8^, much less is known about PCP in the mammalian retina.

A classic model of PCP in vertebrates is the mechanosensory hair cells of the inner ear, which display a V-shaped bundle of mechanosensitive stereocilia oriented in a coordinated manner across the tissue to optimize sound detection^9,10^. Critical to establishing this PCP is the migration of the primary kinocilium from the center of the apical surface to the lateral edge. We and others have reported that relocalization of the kinocilium requires the adaptor protein Inscuteable, the G-protein signaling modulator 2 (GPSM2, also known as LGN) and the heterotrimeric G protein Gαi^11,12^, a protein complex also known for its role in regulating mitotic spindle orientation during asymmetric cell divisions^13–16^. Analogous to the hair cell kinocilium, the connecting cilium of photoreceptors is thought to be anchored eccentrically by the basal body at the edge of the IS. This arrangement constitutes a possible planar asymmetry in photoreceptors. However, it remains unclear whether connecting cilia are coordinately oriented to establish PCP in the mammalian retina. An interesting parallel can be made between the G-protein signaling regulating kinocilium placement in hair cells and the phototransduction cascade, which also relies on a heterotrimeric G-protein. When photons hit the OS disk membranes, the 11-*cis* retinal chromophore isomerizes to all- *trans*, which serves as the ligand for the G-protein coupled receptor opsin. This initiates a biochemical cascade that enables the association of opsin with transducin (Gαt), a heterotrimeric G-protein related to the Gαi family, which in turn activates the cyclic GMP phosphodiesterase controlling cGMP-gated ion channels at the photoreceptor plasma membrane^8,17,18^. Of note, GPSM2 is the only guanine dissociation inhibitor (GDI) protein expressed in mouse photoreceptors, and it was found to interact with the alpha subunit of rod transducin (Gαt1) in a GDP-bound state^19–21^.

Based on these observations, we hypothesized that light might trigger a G-protein signaling cascade modulated by GPSM2, which would lead to the asymmetric positioning of connecting cilia and establishment of PCP in photoreceptors of the mouse retina. Using high-resolution 3D reconstructions of photoreceptors, dark-rearing, mouse genetics, co-immunoprecipitation, and ultrastructure expansion microscopy, we provide evidence supporting this hypothesis.

## RESULTS

### Cone photoreceptors exhibit planar cell polarity in the mammalian retina

Classically, the photoreceptor connecting cilium is presented as eccentrically anchored at the surface of the IS (Fig. 1A), but whether this is a general feature has not been explored in detail. To begin to tackle this question, we investigated the ultrastructure of the mouse retina in 3D using <u>F</u>ocused Ion <u>B</u>eam <u>S</u>canning <u>E</u>lectron <u>M</u>icroscopy (FIBSEM) to obtain hundreds of serial images in the region of the connecting cilia. Transverse and en face milling were collected in the dorsal region of two independent adult mouse retinas (Fig. 1B, C, C’). Computational segmentation of the different sub-compartments of photoreceptors (IS, OS, basal body and connecting cilium) was carried out for each individual rod and cone identified in the volume of acquisition (Fig. 1D; Fig. S1A; Movie S1). By analyzing the relative distance of the basal body to the center of the IS at the surface of each photoreceptor, we discovered that the basal body is systematically located next to the edge of the IS in both rods and cones (Fig. 1C, C’, E), confirming the general assumption that the cilium of photoreceptor cells is intrinsically polarized (i.e. eccentric) relative to the plane of the retina.

**Figure 1.**
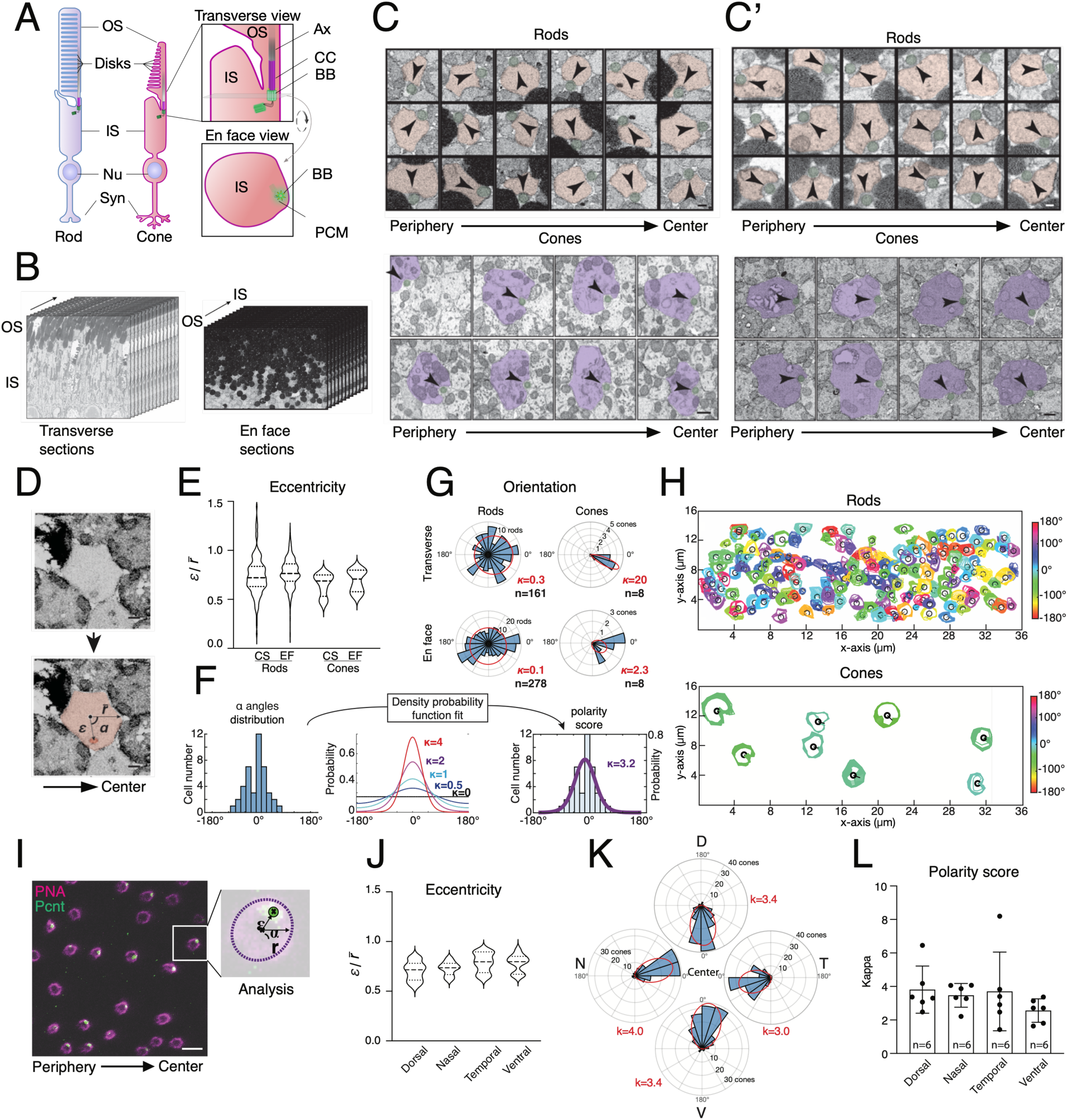
Cones are planar polarized in the adult mouse retina. (**A**) Schematic of photoreceptors basic structure in transverse section. Detail of the connective cilium region (Right panel) and top view of the sliced plane (grey) in the basal body region is shown below. OS: Outer segment, IS: Inner segment, Nu: Nucleus, Syn: Synapse, Ax: Axoneme, CC: Connective cilium, BB: Basal body, PCM: Pericentriolar material. **(B)** Transverse (Left; 628 images, 25nm step) and en face (Right; 914 images, 23nm step) FIBSEM acquisitions in the dorsal region from P42 and P40 mouse retina, respectively. (**C,C’**) High magnification en face views of different rods (Upper panel) and cones (Lower panel) in the connecting cilium region from transverse (C) and en face (C’) FIBSEM acquisitions. Orientation of the cilium (green) is indicated relative to the center of rod (red) and cone (purple) IS by an arrowhead. **(D)** Method for measurement of eccentricity and orientation of the basal body. Basal body (dark red) and IS (light red) are segmented and eccentricity is measured by the ratio of the distance *ε* between centers of the IS and of the basal body and the mean radius r̄ of cone IS. The orientation ɑ is defined by the angle between the axis formed between the centers of the IS and of the basal body and the center-to-periphery axis, using the optic nerve head as a reference for the center of the retina. **(E)** Violin plots of rod and cone basal body eccentricities. CS: cross section (n=164 rods; 8 cones), EF: en face (n=280 rods; 8 cones). **(F)** Analysis workflow of basal body orientations. For each photoreceptor cell type, distribution of α-angle values is fitted to density probability distribution function in a polar referential, resulting in generation of a polarity score (**κ**, kappa). See Experimental Procedures for details. **(G)** Polar histograms of basal body orientations (blue bins) and the associated density probability distribution fit (red ellipse). **(H)** En face view of rod (Top) and cone (Bottom) IS membranes (colored) and basal body (black circle) color-coded for the associated α value obtained in the transverse acquisition. **(I)** Optical slice (confocal) of an adult mouse retina flat mount stained for peanut agglutinin lectin (PNA, magenta) and pericentrin (green). Basal body eccentricity and orientation were measured using the method shown in (D). **(J)** Violin plot of cone basal body eccentricity in different regions of the adult mouse retina by confocal analysis (n= 40 cones from 3 different adult retinas). **(K)** Polar histograms of basal body orientations (blue bins) from different regions of P40 to P60. Dorsal (D); Nasal (N); Temporal (T); Ventral (V). One-way ANOVA: p=0.12. **(L)** Histogram of cone polarity scores (**κ**, kappa) in different regions of the adult mouse retina. Graph shows mean “s.d.; n indicates biological replicates. Total number of cones analyzed were 537 dorsal, 664 nasal, 675 temporal, 628 ventral (see also Fig. S1E). One-way ANOVA: p=0.45. Scale bars: 300nm (C, C’ Upper panels, D), 500nm (C, C’ Lower panel), 10 μm (I).

We next investigated whether this cellular asymmetry may be coordinated across the plane of the retina. To do this, we measured the planar orientation of basal bodies in individual rod and cone photoreceptors, relative to the periphery-to-center centripetal axis of the retina. To help quantify the degree of possible planar polarity, the cumulative distribution of basal body orientations was fitted to a probability distribution function in a polar referential (Fig. 1F; Fig. S1B), generating a polarity score (*κ*, kappa) for each set of orientations, with 2/κ representing the 95% confidence interval in radians and a preferred orientation (*μ*, mu), as previously described^22^. While a kappa value of 0 indicates even distribution of orientations across the circle (i.e. no polarity), kappa values higher than 1 indicate that 95% of orientations encompass less than half of the circle (i.e. polarized). We found that basal body orientations are random among the rod population in both acquisitions (Fig. 1G, H; Fig. S1C). In contrast, we found that basal bodies of cones are consistently located on the side of the IS that is closest to the center of the retina (Fig. 1G, H; Fig. S1D).

Although robust, the coordinated orientation of basal bodies could only be studied for a small region and limited number of cones by FIBSEM, as the mouse retina is rod-dominated. To determine whether cones exhibit PCP in all regions, we used a method that enables a higher throughput of measurements by staining cone IS with peanut agglutinin lectin (PNA) and basal bodies with pericentrin (Pcnt). We then measured basal body eccentricities and orientations on confocal z-stacks of retinal flat -mounts using the same method as in the FIBSEM experiments (Fig. 1I). We found that independent of their location across the retina, cones consistently display eccentric basal bodies located on the side of the cell facing the center of the retina (Fig. 1 J-L; Fig. S1E). Consistent with this observation, we found that cones expressing exclusively one of the two cone opsins also exhibit PCP (Fig. S2A-D).Taken together, these results reveal the intrinsic planar polarity of the basal body in all photoreceptor cells and uncover a tissue-wide PCP of cones (cone PCP) within the mammalian retina.

### Planar polarization of cone basal bodies correlates with the time of eye opening

Cones are amongst the earliest-born cell types in the retina, with peak genesis around embryonic day (E)14.5. Cones OS, however, only starts to grow from the connecting cilium around postnatal day (P)8^23^. To identify when PCP emerges during cone morphogenesis, we first measured basal body orientations at different developmental stages using confocal acquisitions of immunostained retinal flat mounts (Fig. 2A; Fig. S3A). We found that basal bodies are centrally located at the surface of cone IS before P5, and move to an eccentric position between P5 and P8 (Fig. 2A, B; Fig. S3A). At P12, eccentric basal bodies do not yet display a coordinated planar orientation (Fig. 2A, C; Fig. S3B). By P14, however, we observed polarized orientation toward the center of the retina, which is further refined at P16 and remains stable at P30 (Fig. 2A, C; Fig. S3C). These results indicate that cone basal body orientations become coordinated in the plane of the retina around the time of eye opening (P12-P14).

**Figure 2.**
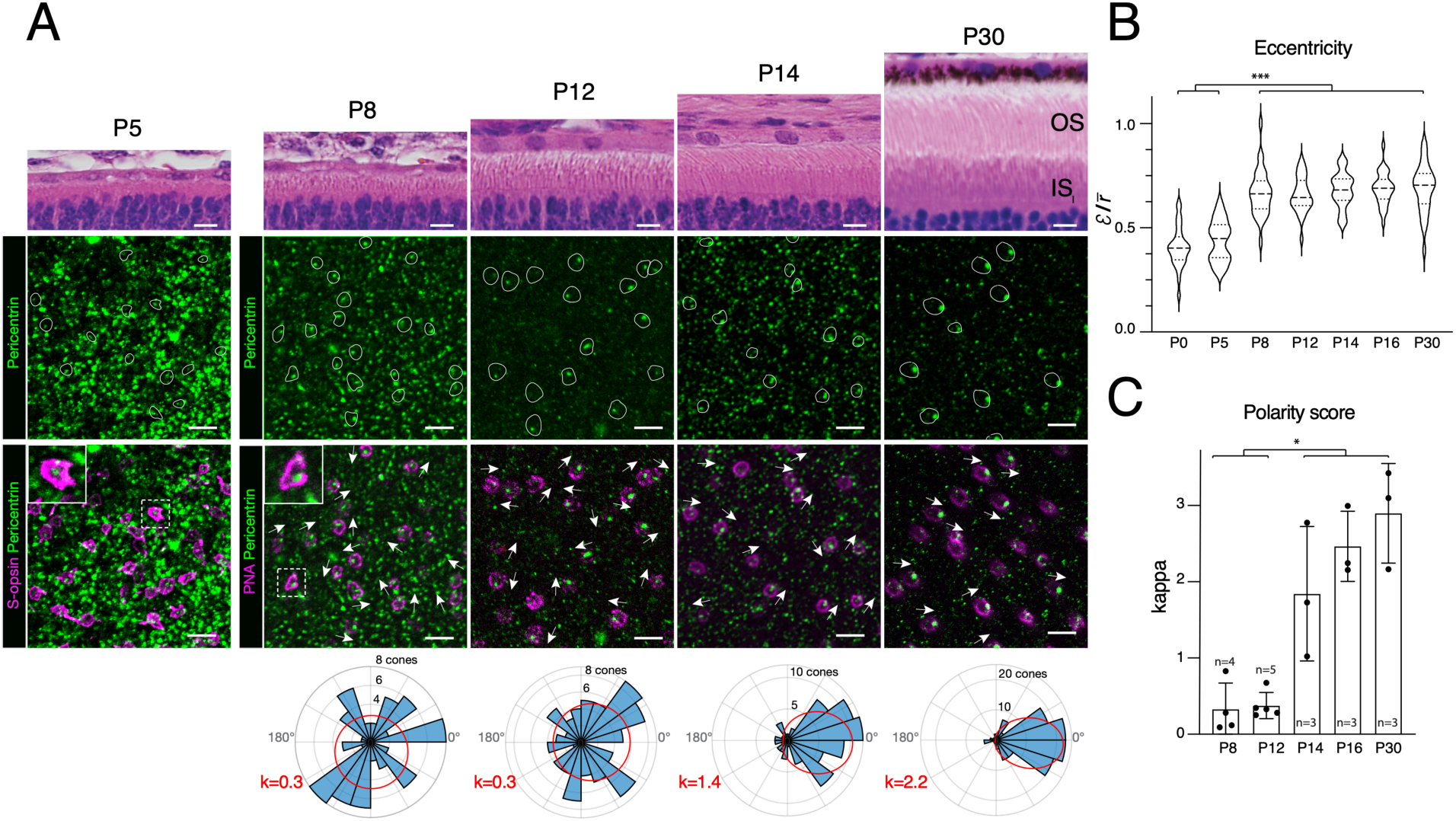
Cone PCP is established by the second postnatal week. **(A)** Hematoxylin/eosin staining of mouse retinal transverse sections at different stages, as indicated, and associated confocal acquisitions of flat mounts of the contralateral retina stained for pericentrin (green), S-opsin (Magenta, P5), or peanut agglutinin (PNA, magenta). Cones are outlined by a white line and basal body orientation is shown by an arrow in the bottom panels. At P5, basal bodies are centrally located while it is eccentric at P8 (zoomed-in insets). For each stage beyond P8, a representative rose plot of basal body orientation (α- angles distribution) and associated kappa value is shown in the bottom row. **(B)** Violin plots of cone basal body eccentricity (n=3 animals per condition, 360 cells counted for each condition). One way ANOVA: p<0.0001. *Post hoc* Tukey’s multi-comparison test: P0, P5 *vs* P8, P14, P16 or P30: p<0.0001(***). **(C)** Histogram of cone polarity scores at different stages. Graph shows mean “s.d.; n indicates biological replicates. Total number of cones measured were: 339 at P0, 299 at P5, 355 at P8, 385 at P12, 1127 at P14, 1389 at P16, and 1314 at P30. One-way ANOVA: p<0.0001. *Post hoc* Tukey’s multi-comparison test: P8, P12 *vs* P14, P16 or P30: p<0.02(*). Scale bars: 10 μm.

### The establishment of cone PCP requires light during a critical period of development

In tissues where PCP has been extensively studied, the Wnt Planar Cell Polarity pathway typically serves as the key regulatory mechanism^1,9,24^. Thus, we next asked whether cone PCP is perturbed in two classical mouse mutants affecting core PCP pathway components: Van Gogh-like protein 2^24^ (VANGL2) and cadherin EGF LAG seven-pass G-type receptor 1^25–27^ (CELSR1). Surprisingly, our analysis of cone PCP in these mouse mutants revealed no discernible effect on cone planar polarization or basal body eccentricity (Fig. 3A-H; Fig. S4A-D), suggesting that an alternative mechanism is at play to establish cone PCP.

**Figure 3.**
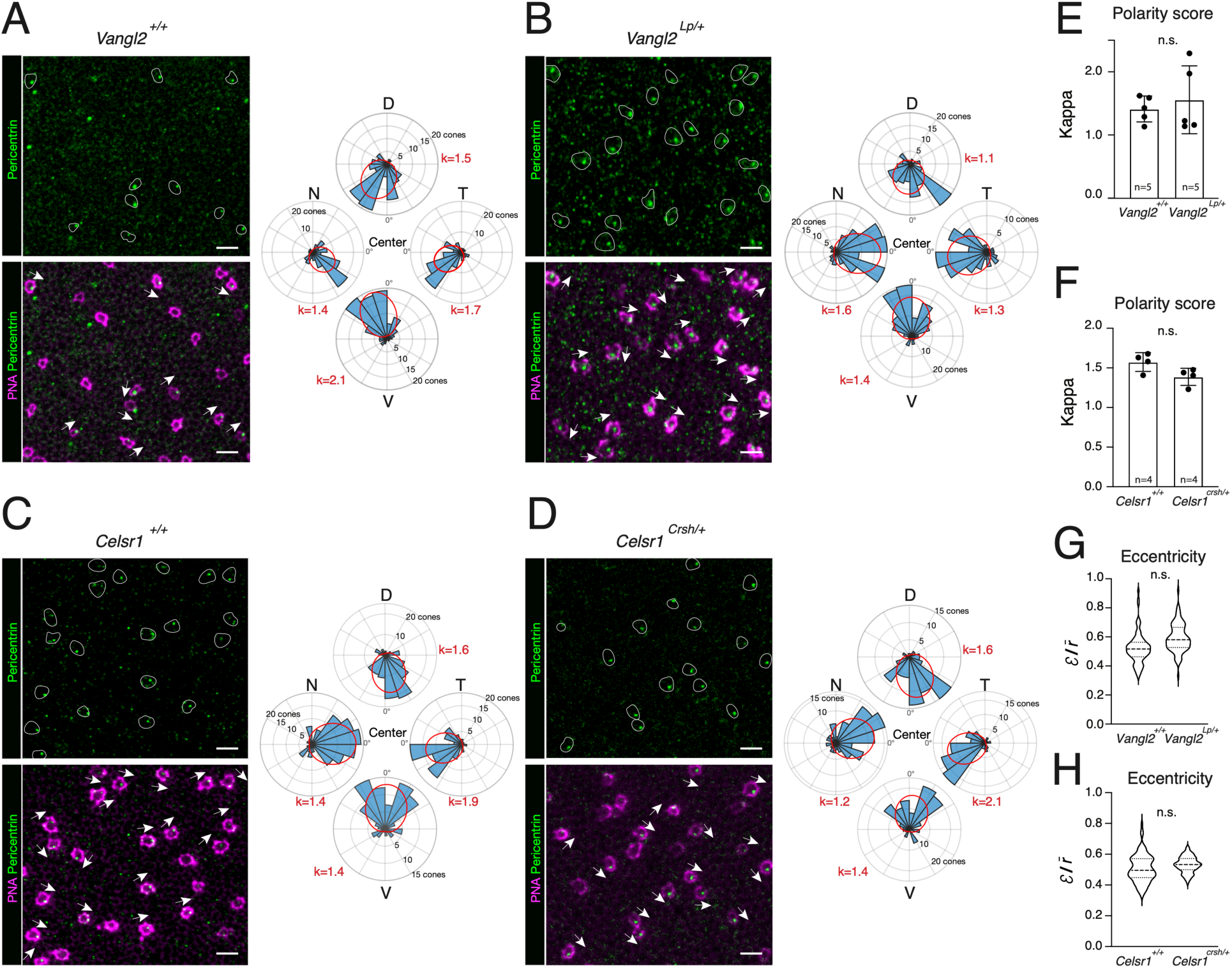
Canonical PCP pathway is not required for cone planar polarization. **(A-D)** Confocal acquisitions of *Vangl2^+l+^* (A), *Vangl2^Lp/+^*(B), *Celsr1^+l+^* (C) and *Celsr1^Crsh/+^* (D) retina flat mounts stained for pericentrin (Pcnt, green) and peanut agglutinin (PNA, magenta). Representative examples of cones (outlined) with various basal body orientations (arrows) are shown. For each genotype, a representative rose plot of basal body orientations (α-angles distribution) and the associated kappa value is shown. **(E-F)** Histograms of cone polarity scores in *Vangl2^+l+^* and *Vangl2^Lp/+^*mice (E), and in *Celsr1^+l+^* and *Celsr1^Crsh/+^* mice (F). Total number of cones analyzed: *Vangl2^+l+^* = 2108, *Vangl2^Lp/+^* = 2384, *Celsr1^+l+^* = 1824, *Celsr1^Crsh/+^*= 1586 cones. Two-tailed unpaired t-test: p= 0.59 (*Vangl2*) and p= 0.90 (*Celsr1*). No statistical difference (n.s.). **(G-H)** Violin plots of cone basal body eccentricity for Vangl2 (G) and Celsr1 (H) genotypes (n=3 animals, 360 cells counted for each genotype). Nested unpaired t-test: p=0.43 (*Vangl2*) and P=0.69 (*Celsr1*). No statistical difference (n.s.). Scale bars: 10 μm.

As cone PCP is established concomitantly with eyelid opening, we next hypothesized that light might be involved. To test this idea, we raised mice in constant darkness from E12.5 and studied cone PCP at P30. Remarkably, light deprivation prevented the emergence of cone PCP, without affecting the eccentric localization of basal bodies (Fig. 4A, E, F; Fig. S5A), showing that light is required for the coordinated arrangement of cilia across the retina plane (cone PCP), but not their lateralization in each cell (intrinsic PCP). To determine whether light is required during a critical window, we raised mice in normal 12h:12h light-dark cycles until P9, P12 or P16, and then switched them to constant darkness until P30. While basal bodies were randomly oriented in mice reared in the dark from P9 (κ=0.6 !0.1; Fig. 4E; Fig. S5A-C) and P12 (κ=0.7!0.1; Fig. 4B, E; Fig. S5A-C), they were planar polarized in mice reared in the dark from P16 (κ=1!0.1; Fig. 4C, E; Fig. S5A-C), although not to the same extent as control mice that were never dark-reared (κ=3.0!0.2; Fig. 4E; Fig. S5A-C).

**Figure 4.**
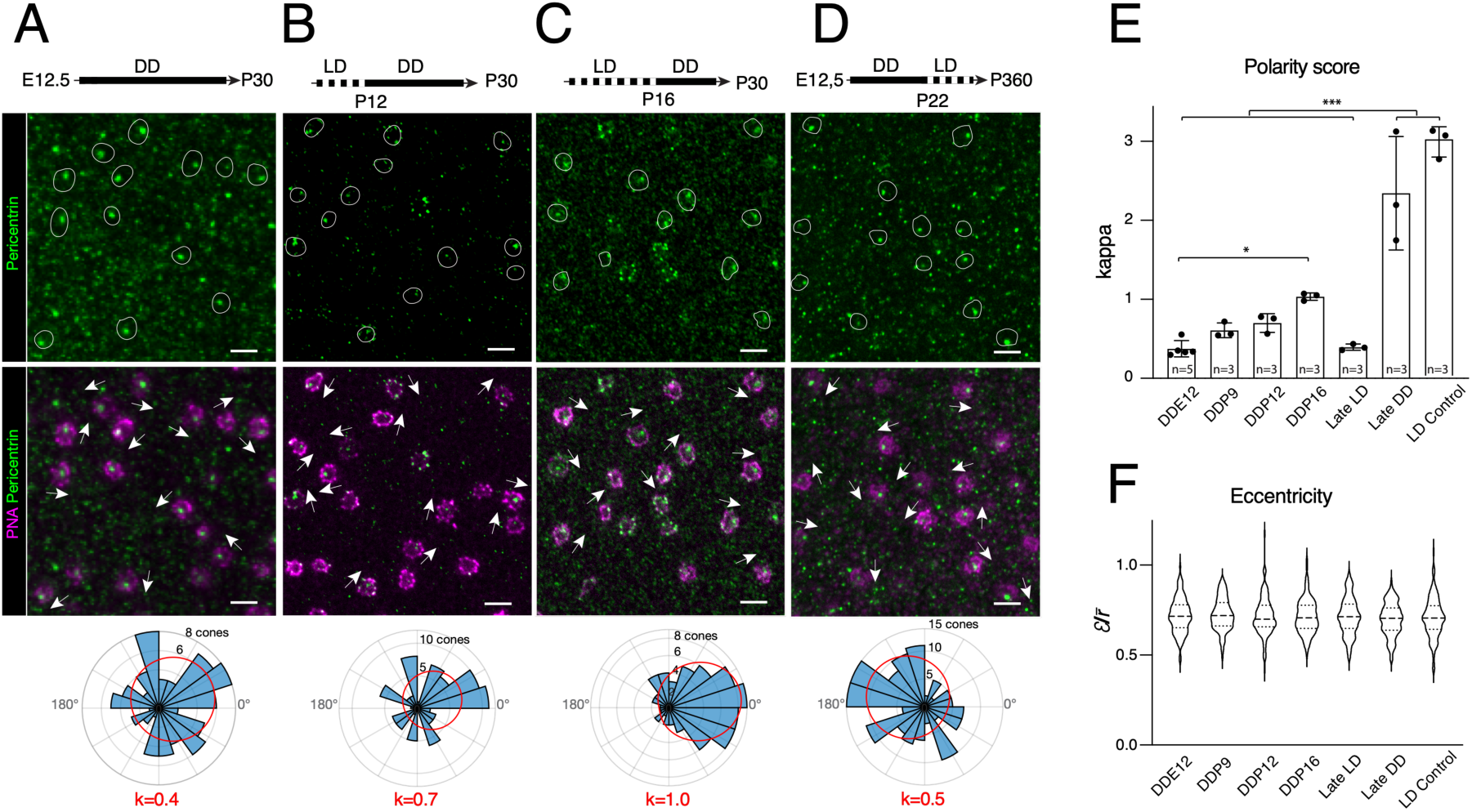
Cone PCP is established during a critical period in development. **(A-D)** Confocal acquisitions of retina flat mounts stained for pericentrin (Pcnt, green) and Peanut agglutinin (PNA, magenta) from dark-reared mice (solid black line) between E12.5 and P30 **(A)**, mice reared in light-dark cycles (dotted line) until P12, then dark-reared until P30 **(B)**, mice reared in light-dark cycle until P16 and dark-reared until P30 **(C),** or mice reared in the dark between E12.5 and P22 then reared in light-dark cycles until sacrifice at P360 **(D)**. Representative examples of cones (outlined) with various basal body orientations (arrows) are shown. For each stage, a representative rose plot of basal body orientations (α-angles distribution) and associated kappa value is shown in the bottom row. **(E)** Histograms of cone polarity scores in mice subjected to different light environments. LD: light:dark cycles, DD: dark:dark cycles, DDE12: dark-reared from E12.5 and until P30, DDP9: dark-reared from P9 and until P30, DDP12: dark-reared from P12 and until P30, DDP16: dark-reared from P16 and until P30, Late LD: dark-reared between E12.5 and P22, then reared on a light-dark cycle until P360, Late DD: mice dark-reared from P120 and until P150, LD control: P30 mice reared in light-dark cycles. Graph shows mean “s.d.; n indicates biological replicates. Total number of cones analyzed: DDE12 = 3417, DDP9 = 1663, DDP12 = 1070, DDP16 = 886, Late LD n=1309 cones, Late DD n= 1077, LD control n=1224 cones. One-way ANOVA: p<0.0001. *Post hoc* Tukey’s multi-comparison test: DDE12.5 *vs* DDP16 p= 0.03(*); DDE12.5, DDP9, DDP12, DDP16, Late LD *vs* Late DD or LD control: p <0.001(***). **(F)** Violin plots of cone basal body eccentricity (n=3 animals, 360 cells counted for each condition). One-way ANOVA: p=0.45. Scale bars: 10 μm.

Importantly, rearing conditions impacted cone PCP in all retinal quadrants homogeneously (Fig. S5B), indicating that cones across the entire mouse retina require light to polarize in the retinal plane. Together, these results show that a light- dependent mechanism is required during a critical period between P12 and P16 to coordinate cone tissue PCP, but not lateralization of cilia (Fig. 4F).

Finally, we wanted to determine whether the disrupted cone PCP observed after dark- rearing could be restored by exposing the mice to light after the critical period. We dark-reared mice from E12.5 until P22, and then switched them to normal light-dark cycles until P360. Despite being exposed to normal light patterns for almost a year after P22, we found that cone basal bodies still exhibit random orientations in these animals (Fig. 4D, E; Fig. S5A-C). Conversely, cone PCP was not disrupted by switching adult mice raised in light-dark cycles to constant darkness for a month (Fig. 4E; Fig. S5A-C). These results show that exposure to light is not sufficient to correct cone PCP defects observed in animals reared in darkness during the critical period, and further suggest that light is not required to maintain cone PCP past the critical window.

### Cone transducin is required to establish cone PCP

The above data indicate that light controls cone PCP, but whether it acts directly on cones, or indirectly through rods remained unclear. To clarify the cellular origin of the light-dependent polarization, we investigated cone PCP in mouse mutants of *Gnat1* and *Gnat*2, which code for the transducin alpha subunits of rods (Gɑt1 protein) and cones (Gɑt2 protein), respectively. Importantly, while rod and cone photosensitivity is abolished in *Gnat1^null/null^* mice and *Gnat2^null/null^* mice, respectively, the general anatomy of the retina and morphology of photoreceptors are unaffected^28,29^. When cone photosensitivity was blocked without altering rod photosensitivity (*Gnat2 ^null/null^*), we found that cone PCP is disrupted without a change in basal body eccentricity (Fig. 5A, B, E, G; Fig. S6A, B). Similarly, when photosensitivity of both rods and cones was blocked (*Gnat1^null/null^*; *Gnat2^null/null^)*, we observed a loss of cone PCP, again without affecting basal body localization (Fig. S6A-H). However, when rod photosensitivity was blocked without altering cone photosensitivity (*Gnat1^null/null^*; *Gnat2^+/null^* mice), we found normal cone PCP and basal body localization (Fig. S6C-H). These results show that the establishment of cone PCP requires photosensitivity in cones, but not rods.

**Figure 5.**
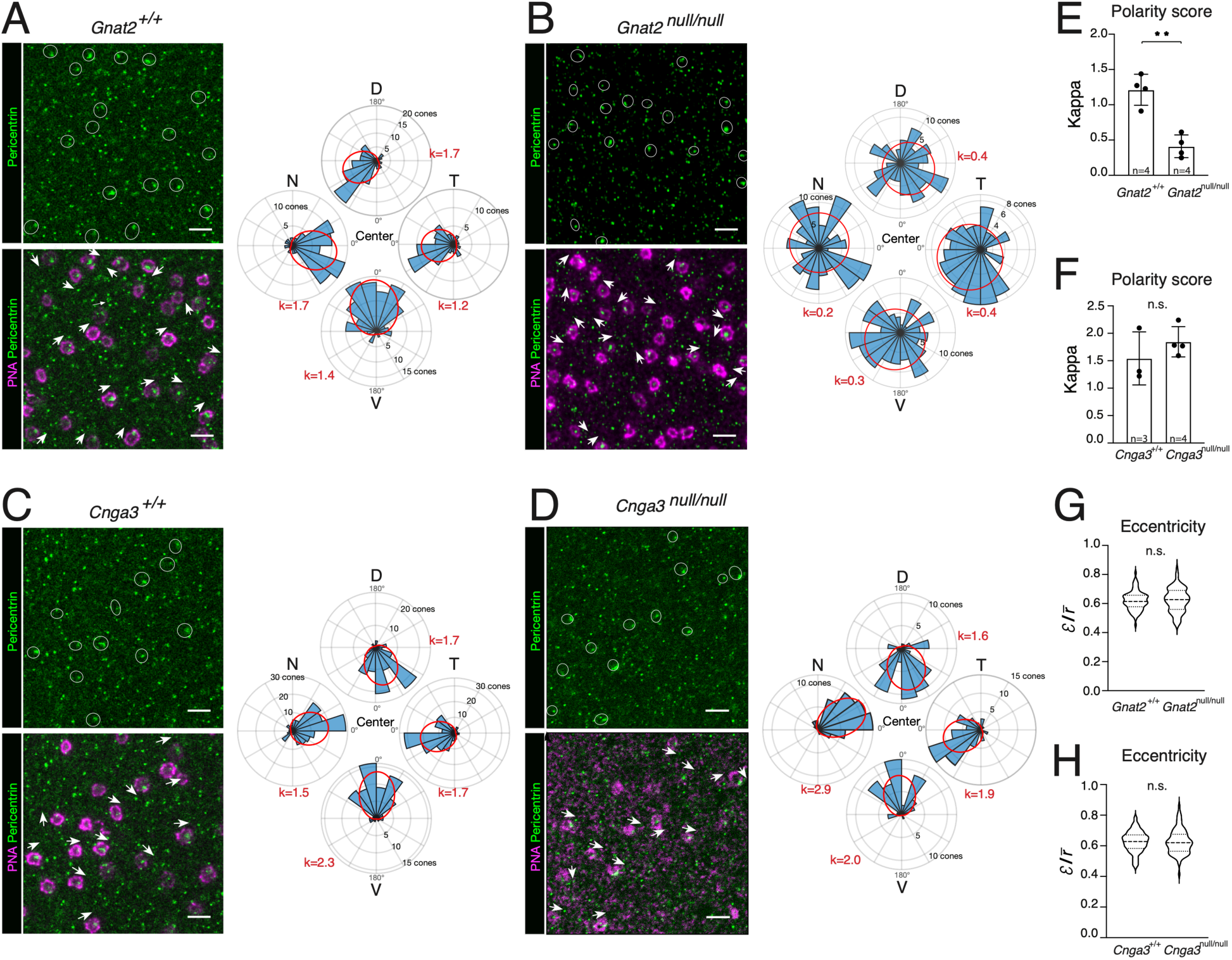
A non-canonical function of transducin underlies cone PCP establishment (A-D) Confocal acquisitions of *Gnat2^+l+^* **(A)**, *Gnat2^null/null^* **(B),** *Cnga3^+l+^* **(C),** and Cnga3*^null/null^* **(D)** retina flat mounts stained for pericentrin (Pcnt, green) and Peanut agglutinin (PNA, magenta). Representative examples of cones (outlined) with various basal body orientations (arrows) are shown. For each genotype, a representative rose plot of basal body orientations (α-angle distribution) and the associated kappa value is shown. **(E-F)** Histograms of cone polarity scores in *Gnat2^+l+^ and Gnat2^null/null^ mice (E), and in Cnga3^+l+^ and Cnga3^null/null^*mice (F). Graph shows mean “s.d.; n indicates biological replicates. Total number of cones analyzed: *Gnat2^+l+^* = 1758, *Gnat2^null/null^* = 1874 cones, *Cnga3^+l+^* = 1525, *Cnga3^null/null^* = 906 cones. Two-tailed unpaired t-test: p=0.001 (\*\**Gnat2*) and p=0.34 (*Cnga3*). **(G-H)** Violin plots of cone basal body eccentricity for *Gnat2* (G) and *Cnga3* (H) genotypes (n=3 animals, 360 cells counted for each genotype). Nested unpaired t-test: p=0.64 (*Gnat2*) and p=0.98 (*Cnga3*). No statistical difference (n.s.). Scale bars: 10μm.

The first steps of phototransduction involves interaction between opsin and the Gα protein transducin, which leads to the closing of cyclic-nucleotide-gated (Cng) channels and hyperpolarization of photoreceptors. To determine whether the entire phototransduction cascade is required for cone PCP, or just the initial steps, we first investigated mouse mutants of *Cnga3*, in which the initial steps of the phototransduction cascade take place normally, but the light-dependent changes in cone cGMP levels are not translated into electrical activity. Strikingly, in contrast to the Gnat2^null/null^ mice, we observed normal cone PCP in Cnga3^null/null^ mice (Fig. 5C, D, F, H; Fig. S6I, J). These results indicate that the initial photosensitive G-protein activity in cones is required to promote PCP, but the resulting ion channel closure and hyperpolarization is not necessary, thereby identifying a non-canonical role for cone transducin in retinal morphogenesis.

### GPSM2 interacts with transducin to establish cone PCP

We next investigated how light detection in cones could lead to the coordinated arrangement of basal bodies within the plane of the retina. Of note, cone transducin (Gαt2) is encoded by the *Gnat2* gene, which arose through duplication of an ancestral *Gnai* gene belonging to a family of heterotrimeric G-proteins, which are regulated by GPSM2 in various developmental contexts^16,30,31^. Thus, we hypothesized that GPSM2 might similarly regulate the activity of cone transducin to elicit cone PCP. To investigate this question, we first stained mouse retinas for cone transducin and GPSM2 during the critical window of cone PCP establishment, using a GPSM2 antibody that we validated on *Gpsm2*^-/-^ samples by western blot and immunostaining (Fig. S7A, B). We found that cone transducin and GPSM2 both localize in the IS of cones between P8 and P14 (Fig. 6A). Furthermore, we found that GPSM2 co- immunoprecipitate with Gαt2 when both proteins are expressed in cultured cells (Fig. S7C) and in retinal extracts (Fig. 6B), consistent with the hypothesis that they interact and function together to regulate cone PCP. Using ultrastructural expansion microscopy^32^, we found that GPSM2 is asymmetrically localized in a stalk along the apico-basal axis of the IS next to the basal body (Fig. 6C-D; Fig. S8A, B). Moreover, by studying en face views, we found that GPSM2 systematically localized on the same side in neighboring cones (Fig. 6E; Fig. S8A), suggesting planar polarization. Importantly, in the basal body region, we observed that GPSM2 polarizes in the retinal plane at P14 in cones of mice reared in normal light/dark cycles (Fig. 6E-G; Fig. S8C), whereas this GPSM2 polarization was lost in mice reared in constant darkness from P5 (Fig. 6E-G; Fig. S8C). Thus, we conclude that light is required to polarize GPSM2 in the plane of the retina.

**Figure 6.**
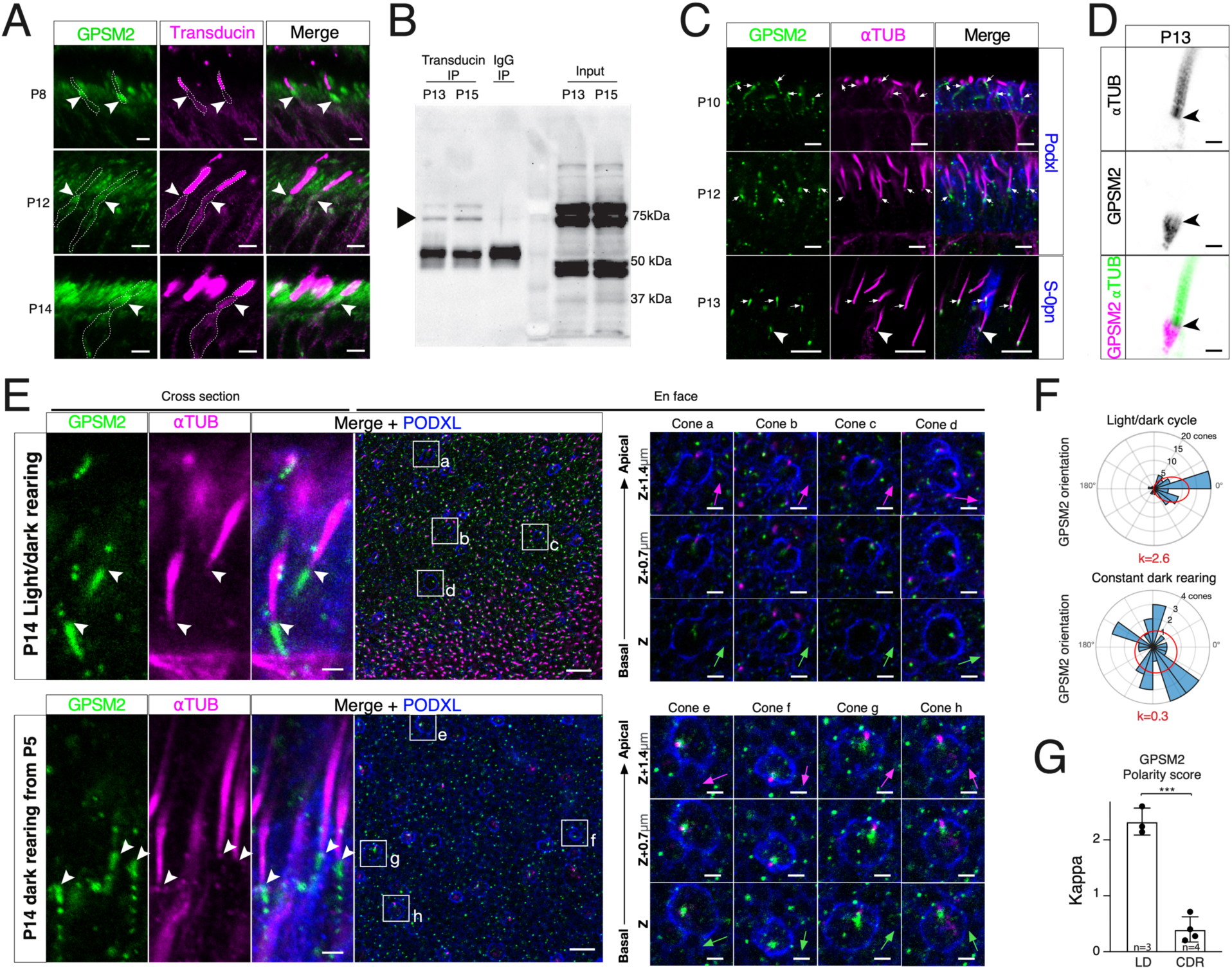
GPSM2 interacts with cone transducin and is polarized by light in the plane of the retina. **(A)** Confocal acquisitions o*f* mouse retina sections stained for GPSM2 (green) and cone transducin (magenta) at different developmental stages, as indicated. Arrowheads point to the cone IS and the dotted line delineate cone contour. **(B)** Co-immunoprecipitation (co-IP) of cone transducin and GPSM2 from mouse retina protein extracts at P13 and P15. Rabbit IgG (IgG) were used as IP controls at P13. 10% input form P13 and P15 whole retina protein extracts are shown. GPSM2 specific band migrates at ∼75kDa (arrowhead, see also Figure S7). **(C)** Confocal acquisitions of mouse retina at different stages after ultrastructure expansion, stained for acetylated tubulin (magenta), GPSM2 (green) and the cone IS marker PODXL^33^ (blue). Arrows and arrowhead point to the distal GPSM2 in the basal body region of rods and cones, respectively. **(D)** Confocal acquisitions of a P13 mouse photoreceptor after ultrastructure expansion stained for acetylated tubulin (αTUB, green) and GPSM2 (magenta). GPSM2 form processes contacting the base of the proximal region of the basal body (arrowhead). **(E)** Confocal acquisitions of expanded P14 mouse retina exposed to normal light conditions (Upper panels) or dark-reared from P5 (Lower panels) and stained for GPSM2 (green) and acetylated tubulin (magenta). Left panel: acquisition in the transverse orientation. Arrowhead point to the distal GPSM2 in the basal body region. Right panels: acquisitions from en face orientation. For each condition, cone distal IS are shown zoomed-in at 3 different z-stack optic slice (step of 0.7um) at the GPSM2/basal body transition. **(F)** For each condition, a representative rose plot of GPSM2 orientations (angles distribution) and the associated kappa value is shown. **(G)** Histogram of GPSM2 orientations polarity scores in cones of P14 mice reared in normal Light/dark cycles conditions (LD) or in constant darkness from P5 (CDR). Graph shows mean “s.d.; n indicates biological replicates. Total number of cones analyzed: LD = 100, CDR = 111 cones. Two-tailed unpaired t-test: p=0.0001(***). Scale bars: 10μm (A, E), 5μm (C), 1μm (E zoom-in), 500nm (D).

The above results suggest that the asymmetrically localized GPSM2 interacts with cone transducin to control cone PCP. To investigate this possibility, we studied cone basal body orientation in *Gpsm2^-/-^* mice. While cone PCP is normal in *Gpsm2*^+/-^ mice (Fig. S9A, C, E, F), we found that it is disrupted in *Gpsm2^-/-^*mice (Fig. S9B, C, E, F). This finding was further confirmed using FIBSEM analysis on the dorsal region of a *Gpsm2^-l-^* mouse (Fig. S9I-M). To determine whether *Gpsm2* is required cell-autonomously in cones to promote PCP, we generated a cone-specific conditional KO of *Gpsm2* by crossing a *Gpsm2^fl/fl^* mouse line to the Opn1LW-Cre mouse line, which expresses Cre specifically in cone photoreceptors from P10^34^, prior to the opening of the critical period for cone PCP establishment. Whereas we detected normal cone PCP in the Opn1LWCre+; *Gpsm2^fl/+^* control mice (Fig. 7A, C; Fig. S9G, H), it was disrupted in the Opn1LWCre+; *Gpsm2^fl/fl^* cKO mice (Fig. 7B, C; Fig. S9G, H), demonstrating that GPSM2 function in cones is required to promote PCP. In contrast to previous findings in cochlear hair cells^11,12,35^, however, the loss of cone PCP in *Gpsm2^-/-^* or *Gpsm2* cKO was not accompanied by a disruption of the eccentric localization of cone basal bodies (Fig. 7D; Fig. S9D). Together, these results identify a molecular pathway downstream of light involving both transducin and GPSM2, which is necessary to elicit cone PCP in the mouse retina.

**Figure 7.**
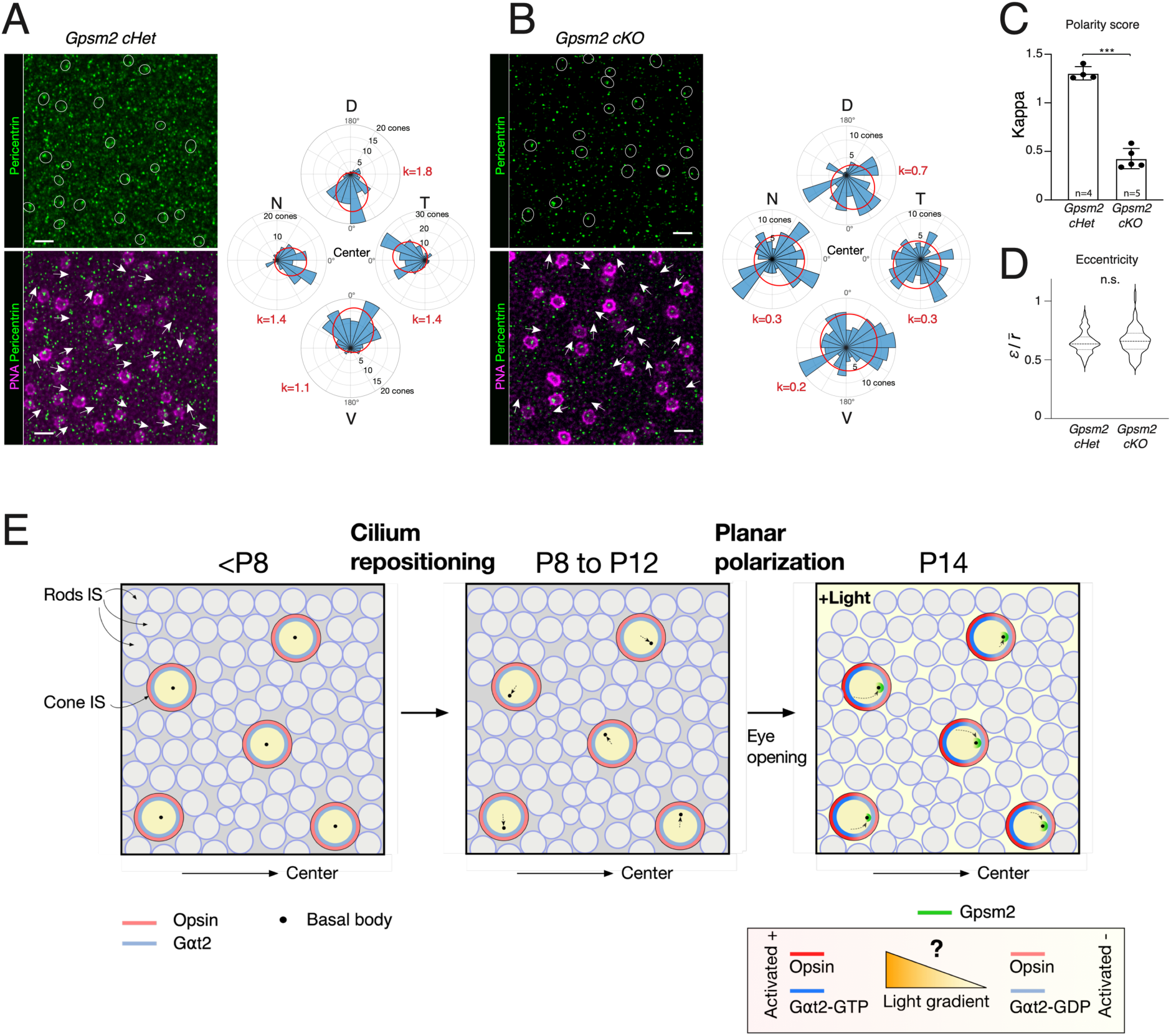
GPSM2 is required for cone PCP. **(A, B)** Confocal acquisitions of *Gpsm2* conditional heterozygote (cHet) **(A)** and conditional knock-out (cKO) **(B)** retina flat mounts stained for pericentrin (Pcnt, green) and peanut agglutinin (PNA, magenta). Representative examples of cones (outlined) with various basal body orientations (arrows) are shown. For each genotype, a representative rose plot of basal body orientations (α-angles distribution) and the associated kappa value is shown. **(C)** Histogram of cone polarity scores in *Gpsm2 cHet and Gpsm2* cKO mice. Graph shows mean “s.d. n indicates biological replicates. Total number of cones analyzed: *Gpsm2 cHet* = 2008, *Gpsm2 cKO* = 2799 cones. Dorsal (D); Nasal (N); Temporal (T); Ventral (V). Two-tailed unpaired t-test p<0.0001 (***). **(D)** Violin plots of cone basal body eccentricity (n=3 animals, 360 cells counted for each genotype). Two-tailed unpaired nested t-test: p= 0.44. No statistical difference (n.s.). (**E**) Model depicting how cone cilia polarize over time. Initially central before P8 (left panel), basal bodies reposition between P8 and P12 to become eccentric and contiguous to the IS membrane (intrinsic PCP), but do not display any preferred orientations in the retinal plane (central panel). Between P12 and P14, basal body positions become coordinated across the tissue plane to sit on the side of the IS closer to the center of the retina (tissue PCP, right panel). This last step requires light upstream of both cone transducin and GPSM2. A speculative mechanism integrating all 3 factors is also depicted (below Right panel). Arrows depict the early lateralization shift (Central panel) and subsequent relocalization of the cilium (Right panel). See Discussion for details. Scale bars: 10 μm.

## DISCUSSION

Extensive studies in various sensory tissues have established that PCP is primarily controlled by a canonical core PCP signaling pathway. This pathway integrates molecular gradients to regulate inter-cellular communication, thereby coordinating the orientation of polarized cellular structures^1,3^. In this study, we uncover PCP organization in all cone subtypes of the mouse retina and show that it is established through a non-canonical PCP pathway triggered by light (Fig. 7E).

In several aspects, the establishment of cone PCP is reminiscent of the mechanism observed in hair cells of the inner ear. In both tissues, PCP is established through a two-step process: lateralization of the cilia followed by their coordinated planar polarization within the tissue (Fig. 7E). This morphogenetic sequence, as well as the contribution of a G-protein pathway coupled to GPSM2 activity, closely resembles the process responsible for PCP establishment in the ear^11,12,36^. However, the specific role of GPSM2 in this process, in addition to the role of light, differs significantly between the two tissues. While in the inner ear, GPSM2 activity promotes cilium lateralization, in cones, the role of GPSM2 is limited to coordinating cilia positions within the tissue plane. Further studies await to understand this difference, but a possible explanation may lie in the distinct sub-cellular configurations of cilia, particularly the presence of an inner segment in cones that separates basal bodies and tight junctions in distinct planes.

Our discovery raises a fundamental question about the role of light in establishing cone PCP. One possibility is that light acts in a permissive manner to activate an intrinsic pathway that subsequently guides PCP through classical gradients of molecular cues. While this scenario cannot be entirely ruled out, our findings that mutants in the core PCP pathway, which have altered PCP in the inner ear and other organs^24–26^, display normal cone PCP in the retina, weakens this hypothesis. An alternative possibility is that light itself, rather than molecules, serves as the primary instructive cue for cone planar polarization. In such model, exposure to light might trigger specific cellular responses that directly influence the organization and alignment of cone cells in the retina. While we favour the second model, additional work will be necessary to directly address whether light plays a permissive or instructive role in cone PCP.

Molecularly, our data indicate that GPSM2 and cone transducin (Gαt2) interact and are both required to establish cone PCP. In asymmetric cell division, GPSM2 functions as an adaptor protein linking Gαi subunits anchored at the apical cell cortex to the microtubule and the dynein-associated Nuclear mitotic apparatus protein (NUMA), leading to capture of aster microtubules and reorientation of the mitotic spindle^13,15,37^. In a striking example of pleiotropy, we found that the position of the microtubule-based basal bodies in cones is also regulated by GPSM2 and the Gɑi-related protein cone transducin. While GPSM2 and cone transducin constitute the intracellular levers to position the basal body in cones, the mechanisms coordinating basal body positions across the retina remain to be determined. As light is the “ligand” sensed by the photoreceptor specific GPCR opsin, our observations raise the possibility that a gradient of light, rather than molecules, instruct basal body positioning during cone maturation (Fig. 7E). In such a model, cone transducin may constitute the molecular link between light and GPSM2, forming a non-canonical pathway to instruct the spatial localization of cone basal bodies across the retina. This model is grounded on the exclusive interaction of GPSM2 with the alpha subunit of transducin in its GDP conformation^12,38,39^. The interaction of GPSM2 with Gɑi-GDP results in unfolding of GPSM2 structure, allowing for the recruitment of microtubule associated proteins to the Gɑi-GPSM2 complex ^13,15,37^. We therefore postulate that a gradient of light could result in an asymmetric activation of GDP- vs. GTP-loaded transducin, leading to the recruitment of GPSM2 and basal bodies towards the side of lower illumination (Fig. 7E) but, as mentioned above, the possibility that light simply activates a classical PCP pathway cannot be excluded at this time.

By sampling critical information in the environment, sensory organs such as the retina are particularly prone to morphological changes related to their function. While it remains to be determined whether cone PCP in mammals is an adaptation to nocturnal vision, it is worth noting that polarization of cone basal bodies was reported in the zebrafish retina, a diurnal animal with a cone-dominated retina^40^. In zebrafish, red, green, and blue cones harbor an eccentric basal body pointing towards the optic nerve head, as we observe here in mice, while UV-cone and rod PRs do not. Although the mechanisms and role of this polarization in fish were not identified^41^, it was reported to arise at late stages of development, suggesting that light, as we find in mice, may be involved. Recently, cones of the primate retina were shown to exhibit planar polarization, albeit in a bimodal fashion, with the basal body more frequently anchored on the edge of the IS facing either the periphery or the center of the macula^42^. Whether light is also part of the mechanism in primates remains to be determined, but the tropism for cilia towards and away from the macula suggest that it may rely on optical properties rather than a morphogen gradient secreted from the optic nerve head. It will be important to study photoreceptor planar polarity in detail in a wider range of animals to help determine whether cone PCP is the rule, or an exception, across mammals.

Our findings that rod photoreceptors do not exhibit PCP suggest a potential role for PCP in visual acuity, a function predominantly supported by cones. Psychophysical experiments conducted in humans reported that the retina is more sensitive to light entering the center of the pupil than light passing through its periphery and thereby hitting photoreceptors at an angle, a phenomenon referred to as the Stiles-Crawford effect ^43,44^. While this effect is prominent in cones, it is weak in rods^45^, and studies on the optical properties of individual rod and cone PRs have failed to identify a convincing cellular origin of this phenomenon in humans^46,47^. Interestingly, a recent study in mice proposed that the rectilinear orientation of IS and OS of rod photoreceptors, which is instructed by light between P0 and P8, could be involved in the Stiles-Crawford effect^48^. As rods play a minor part in the Stiles-Crawford phenomenon, however, it is likely that cone-specific mechanisms also contribute. *Ex-vivo* experiments suggest that, in contrast to rods, cones do not exhibit a dynamic phototropic realignment of their IS and OS in adult mice^49,50^, suggesting alternative mechanisms. The potential role of cone PCP in the Stiles-Crawford effect is intriguing. We propose that the coordinated orientation of cilia may physically restrict cones to exclusively detect light that is paraxial to the eye optics, filtering out intraocular stray light that deviates from this axis. Our results, showing that cone PCP cannot be reversed by rearing adult mice in the dark, align with psychophysical observations showing that monocular light deprivation fails to alter the Stiles-Crawford effect in adult human subjects^51,52^.The inability to recover PCP beyond the critical period despite exposure to light in dark-reared represents another intriguing discovery. This result contrasts with the permissive critical window identified for the phototropic rectilinear alignment of rod IS in mouse^48^. While further investigations are required to identify the mechanism driving rod IS phototropism, the differing role of transducin could potentially underlie this difference. Based on our findings, it may be possible to test the role of cone PCP in the Stiles-Crawford effect by performing psychophysical experiments on patients bearing a mutation in the *GPSM2* gene, which cause the Chudley-McCullough syndrome, a congenital disorder characterized by hearing loss and various brain anomalies^35,53^.

As many tissues are sensitive to light, our findings raise the possibility that light might regulate PCP in a range of biological contexts^54^, and more broadly influence cell-cell organization. Even during embryo development, recent studies have shown that light penetrates through the womb and can elicit a response in retinal cells^55,56^. Outside the eye, brain regions are naturally sensitive to light^57,58^, raising the exciting possibility that light may generally regulate tissue morphogenesis in the CNS.

## ACKNOWLEDGMENTS

We would like to dedicate this paper to the late Julian Hart Lewis, FRS (LRI, Cancer Research UK), who initially raised the question of photoreceptor PCP in a discussion with M.C. in 2000. That discussion was the spark that eventually led to the start of this project almost 20 years later. We thank J. Barthe, M.-C. Lavallée and C. Dubé for help with the animal colony, S. Terouz for histology, F. Couderc and M. Rondeau for genotyping. We are grateful to S. Hattar (Johns Hopkins) for sharing the Gnat1/Gnat2 mutant mice, B. Tarchini (Jackson Laboratories) and P. Gros (McGill) for sharing the Vangl2^Lp^ mutant mice, D. Devenport and K.A. Little (Princeton University) for sharing the Celsr1^Crsh^ mutant mice, K. Sears, J. Mui, L. Monaghan, L. Mongeon and W. Leelapornpisit at the Facility for Electron Microscopy Research (McGill University) for help in sample processing and data collection by FIBSEM and the FRQS Vision Health Research Network (VHRN) for tissues. This work was funded by grants from the Canadian Institutes of Health Research (FDN-159936 to M.C.; 130376 to C.M.), The Brain Canada/Weston Foundation, and the Natural Sciences and Engineering Research Council (to C. C.). M.H. was funded by a postdoctoral fellowship from the Fondation pour la recherche médicale française (FRM). M.C. is a FRQS Emeritus Research Scholar and holds the IRCM Foundation Gaëtane and Roland Pillenière Chair in Retina biology.

## EXPERIMENTAL PROCEDURES

### Mice

All animal work was carried out in accordance with the Canadian Council on Animal Care guidelines and approved by the IRCM Animal Care Committee. Wild-type C57BL/6J, *Gnat2^Cpfl^*^3^*^/Cpfl^*^3^ (described as *Gnat2^null/null^*in the text), *Cnga3^Cpfl^*^5^*^/cpfl^*^5^ (described as *Cnga3^null/null^* in the text), transgenic *Opn1LWCre+*, and CD1 mice were obtained from The Jackson Laboratories. Retinas from the *Gnat1^irdr/irdr^*; *Gnat2^Cpfl^*^3^*^/Cpfl^*^3^ mice (referred to *Gnat1^null/null^*; *Gnat2^null/null^* in the text) were generously provided by Dr. Samer Hattar (NIMH), the Vangl2^Lp^ mice were generously provided by Dr. Basile Tarchini (Jackson laboratory) and Dr. Philippe Gros (McGill), from the Celsr1^Crsh^ mice were generously provided by Dr. Danele Devenport (Princeton). Germline and conditional *Gpsm2* KO mouse lines were maintained on a mixed C57BL/6J/129S1 background and PCR genotyping for *Gpsm2* and Cre were carried out as previously described^12^. Both male and female animals were used at the indicated developmental and adult time points.

### Immunostainings

After labeling the dorsal pole of the eye with a marker, eyes were enucleated and fixed. Eyes were rinsed 3 times 10 minutes in PBS and dissected to isolate the retina. Four incisions were made to allow flattening of the retina and the dorsal quadrant was additionally incised for identification. Retina flat mounts were then fixed by immersion in 10% trichloroacetic acid (TCA) for 10 minutes at room temperature and permeabilized in blocking solution (0.3% Triton X-100 + 1% bovine serum albumin) overnight at room temperature. Whole-mount primary and secondary antibody incubations were done overnight at room temperature in the blocking solution + 1% thimerosal (Sigma Aldrich, T5125). Samples were washed in PBS overnight between primary and secondary antibody incubations, and between secondary antibody and mounting. After immunostaining, samples were flat mounted in Mowiol (Sigma Aldrich cat#81381). For immunohistochemistry on cross-sections, eyes were fixed by immersion in paraformaldehyde (PFA) 4% for 30 minutes at room temperature, rinsed 3 times 10 minutes, dissected to remove the cornea and the lens, than immersed in 20% sucrose, embedded in 1:1 20% sucrose:OCT (Tissue-Tek), frozen in liquid nitrogen, and stored at −80°C until sectioning. Images were acquired using Leica SP8 confocal epifluorescent or Zeiss LSM700 confocal microscopes using the 63X lens. For histology, eyes were fixed following a previously published protocol (Sun et al., 2015). Briefly, after enucleation, eyes were snapped frozen in propane at dry-ice temperature, immersed in 2.5% acid acetic in methanol and kept 48 hours at -80°C, then progressively warmed to room temperature. Dehydrated retinas were embedded in paraffin, sectioned (5 µm) and stained with hematoxylin and eosin.

### Primary Antibodies

The following primary antibodies were used: rabbit anti-Arr3 (cone arrestin, 1:1000, Millipore Sigma cat#AB15282), rabbit anti-GPSM2 (Abcam cat#AB84571) for western blot (1:500) and immunoprecipitation (4μg/IP), goat anti-GPSM2 (1:500, Life Technology cat#PA5-18646) for immuno-histochemistry, rabbit anti-GPSM2 (1:500, ProteinTech 11608-2-AP), goat anti-PODXL (1:500, R&D systems cat#1556), goat anti-S-opsin (1:2000, Santa Cruz Biotechnology #Cat-SC14363) and mouse anti-acetylated tubulin (1:1000, Sigma-Aldrich cat#T6793) for immunolabeling of expanded retinas. Rabbit anti-Gαt2 (Abcam, cat#ab97501) for immunoprecipitation (4μg/IP) and immuno-histochemistry (1:1000), rabbit anti-PCNT (1 :500, BioLegend cat#923701), rabbit IgG isotype for IP control (4μg/IP; Thermo Scientific cat#02-6102), rabbit anti-FLAG (4μg/IP; Cell signaling Cat#2368P), rabbit anti-GFP (1:2000; Thermo Fisher Scientific Cat#A11122). All secondary antibodies and lectins were diluted 1:1000 in blocking solution + 1% thimerosal. The following secondary antibodies were used: Alexa Fluor (AF)555 donkey anti-rabbit (Abcam, cat#ab150062), AF488 donkey anti-goat (Abcam cat#ab150133), AF647 Donkey anti-Rabbit (Jackson ImmunoResearch cat#711-605-152), AF647-conjugated peanut agglutinin lectin (Molecular probes cat# L-32460). Hoechst was added with the secondary antibody (1μg/mL; ThermoFisher Scientific, cat#H3570).

### Quantification of basal body positions and planar orientation

Measurements were made on confocal z-stacks acquired in the region of photoreceptor IS and OS of TCA-fixed mouse retina flat mount stained for pericentrin, cones (PNA-AF647 and S-opsin for P0 and P5 time points) and DNA (Hoechst). Since immature cones do not exhibit strong PNA staining, S-opsin was used to label cone IS before P8. Stacks were quantified in flat regions located at equal distance between the optic nerve head and the periphery of the retina. ɑ-angle was measured using FIJI (NIH, v2.1.0) as the angle formed between the center of the cone ellipsoid (based on PNA labelling) and the center of pericentrin, with the optic nerve head systematically oriented toward the right of the acquired field (0-degree orientation). While all cones displayed a basal body punctum, its localization varied across different z-planes among cones. This variation explains why some cones do not show basal body puncta in some of the images selected for illustration in the figures (single z-plane). In rare cases where pericentrin labeling resulted in elevated background staining within the IS, we consistently considered the punctum of highest intensity as the basal body for quantification. The orientation of the basal body was measured identically between conditions on a minimum of 30 cones in a flat region to minimize bias linked to cell deformation. Eccentricity (ε) was calculated as the ratio of the distance from the center of the cone ellipsoid to pericentrin and the mean radius (inner periphery) of the PNA-positive shape (r). For comparison of eccentricity, 30 cones from each region (dorsal, nasal, temporal, ventral) of at least 3 independent mice were used between conditions or genotypes except for P0, P5, P8 and P12 retinas in which quantification was made on randomly selected regions from 3 independent mice.

Deformation of the naturally curved retina linked to flat mounting introduces variability in the relative direction of the optic nerve head. To circumvent this caveat, we fitted ɑ- angle distributions to a Von Mises distribution function. The Von Mises distribution function offers a statistical framework for modeling directional data, allowing to objectively quantify the degree of alignment or randomness in observed angle distributions. Fitting this distribution function to the data allows to evaluate the strength and consistency of preferred orientations. A narrow, tall Von Mises curve signifies a high degree of alignment, with angles tightly clustered around a central direction, while a wider, flatter curve suggests a more uniform distribution, indicative of randomness (Fig. S1B). This rigorous approach ensures adherence to academic standards in quantitative analysis across diverse fields, including materials science, ecology, and neuroscience. Utilizing data fitted to a Von Mises distribution function facilitates the derivation of a polarity score (κ) and the direction of the orientation (μ). To get the κ and μ parameter values from the Von Mises distribution, we used the cumulative distribution of angle values and assigned an equal normalized weight to each angle data. We approximated the cumulative probability density function with 1000 modified Bessel functions of order j evaluated at the value κ (see Eq 1) as the model cumulative Von Mises distribution function to fit. We used the least mean square function lsqcurvefit from Matlab (Mathworks) to find the κ and μ parameters under limit constraints of 0.001 < κ < 100 and –π < μ < π which best fit the experimental data.

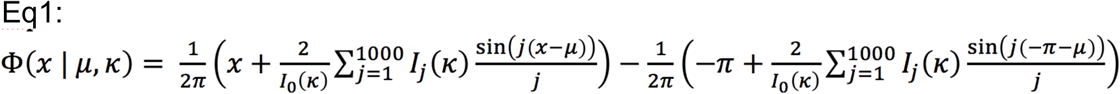

For comparison of polarity scores between retinal regions, mean κ from 6 independent wild type mice (C57B6J) aged between P40 and P60 were used. For comparison of polarity scores between different ages or rearing conditions, the mean κ of the dorsal, nasal, temporal and ventral region from 3 independent wild-type mice (CD1) were used in dark-rearing experiments and for P14, P16 and P30 times points. κ from randomly selected regions from at least 4 independent wild-type mice (CD1) were used for P0, P5, P8 and P12. For comparison of polarity scores genotypes, the mean κ of the dorsal, nasal, temporal and ventral region from at least 3 independent *Vangl2*^Lp/+^, *Vangl2*^+/+^*, Celsr1*^Crsh/+^*, Celsr1*^+/+^, *Cnga3*^null/null^, *Gnat2*^+/+^, *Gnat2*^null/null^, *Gnat1^n^*^ull/null^; *Gnat2^+/^*^null^, *Gnat1^null/n^*^ull^; *Gnat2^nul/n^*^ull^, *Gpsm2^+/-^*, *Gpsm2^-/-^*, Gpsm2 cHet and Gpsm2 cKO mice were used.

### FIBSEM Tomography

Mice were perfused intracardially with 10mL of sodium cacodylate buffer 0.1M, 0.1% CaCl2, pH7.4, followed by 10mL of 1% fixative solution (1% PFA; 1% (vol/vol) glutaraldehyde in sodium cacodylate buffer, and finally with 10mL of a 2% fixative solution (2% PFA; 1% (vol/vol) glutaraldehyde in sodium cacodylate buffer). The dorsal pole was marked, and the eyes were enucleated and immersed overnight in 2% fixative solution + 10% sucrose at 4°C. The 1 mm^3^ of the dorsal retina was dissected and washed 3 times with sodium cacodylate buffer for 30 min, post-fixed in 0.1 M sodium cacodylate buffer containing 1% (wt/vol) OsO4 and 1.5% (wt/vol) potassium ferrocyanide for 2 h at 4 °C. After 3 washes of sodium cacodylate buffer for 30 min, the samples were immersed for 1 hour at 4°C in sodium cacodylate solution and 1% tannic acid. Samples were rinsed 3 times in dH2O for 30 minutes and en bloc stained for one hour with 1% (wt/vol) aqueous uranyl acetate at 4 °C.

Samples were dehydrated in five successive steps of acetone and dH2O [30 to 90% (vol/vol)], each for 15 min at room temperature followed by 100% acetone (3 × 20 min). The samples were incubated with increasing concentrations [30 to 100% (vol/vol)] of low viscosity EPON 812 replacement (Mecalab Limited, Montreal, Canada) over a period of 24 h, and then polymerized at 65 °C for 48 h. Ultrathin sections (70–100 nm) were en face cut from the resin blocks using a Leica Microsystems EM UC7 ultramicrotome (Leica Microsystems, Wetzlar, Germany) with a Diatome diamond knife (Diatome Limited, Nidau, Switzerland) and stained with 1% toluidine blue (Sigma Aldrich cat#T3260) to ensure the quality of the preparation and locate the region of interest (ROI) before data collection by FIBSEM.

The blocks were trimmed with a razor blade to expose the ROI, mounted on fixed 45° pre-tilt SEM stubs, and coated with a 4-nm layer of Pt using a Leica Microsystems EM ACE600 sputter coater (Leica Microsystems, Wetzlar, Germany) to enhance electrical conductivity. Milling of the serial sections and imaging of the block face after each Z-slice was carried out by the Helios Nanolab 660 DualBeam using Auto Slice & View G3 ver 1.2 software (Thermo Fisher Scientific). The sample block was first imaged to determine the orientation of the block face and ion and electron beams. A 2-µm layer of Pt was deposited on the surface of the ROI to protect the resin volume from ion beam damage and correct for stage and/or specimen drift, i.e. orthogonal to the block face of the volume to be milled. Trenches on either side of the ROI were milled to minimize redeposition during the automated milling and imaging. Fiducials were generated for both the ion and electron beam imaging and used to dynamically correct for drift in the x- and y-directions during data collection by applying appropriate SEM beam shifts. Milling was performed at 30 kV with an ion beam current of 9.3 nA, stage tilt of 4°, and a working distance of 4 mm.

At each step, a 20 nm (for cross-section acquisitions) or 23 nm (for en face acquisitions) slice of the block face was removed by the ion beam. Each newly milled block face was imaged with the through-the-lens-detector (TLD) for backscattered electrons at an accelerating voltage of 2 kV, beam current of 0.40 nA, stage tilt of 42°, and a working distance between 2.5 to 3.5 mm. The pixel resolution was 17 nm (cross-section) and 23.6nm (en face) with a dwell time of 10 μs per pixel. Pixel dimensions of the recorded images were 2048 × 1768 pixels (cross-section) and 1536x1034 (en face). Six hundred twenty-eight images (cross-section) or 914 images (en face) were collected from wild-type (C57/Bl6) mouse retina. Seven hundred twenty-eight images (en face) were collected from the Gpsm2^-/-^ mouse retina. Visualization and direct 3D volume rendering of the acquired datasets was done with Amira for Life Sciences software (Thermo Fisher Scientific. Hillsboro, Oregon).

Cone photoreceptors were distinguished from rod photoreceptors based on the difference in IS size and shape of the OS. The photoreceptor IS membrane and basal body were manually segmented over the acquired images (enface) or on images after z-projection (cross-section) for different z planes using the MatLab program (The Mathworks Inc., Natick, MA). Missing in-between z planes were linearly interpolated. For planes where the IS and basal body of the same photoreceptor overlapped, coordinates of the border regions from each object were averaged to generate geometric centers and used to calculate the orientation and eccentricity (see quantification section above).

### Dark Rearing

Timed-pregnant wild-type females (CD1, plug date set as E0.5) were placed in complete and constant darkness from E12.5 onward and pups were either sacrificed at P30 or reared in a light:dark (LD) cycles from P22 onward and sacrifice at P360. Alternatively, wild-type mice (CD1) were placed in complete and constant darkness from P8, P12 or P16 onward and sacrificed at P30. LD controls were maintained on a 12:12 light:dark cycle with ambient light-phase intensity averaging 200 lux.

### Co-immunoprecipitation

Wild-type C57B6J mice were euthanized by CO2 asphyxia, and enucleated. Eyes were dissected individually in cold 1X phosphate-buffered saline (PBS) to isolate the neural retina. Isolated retinas were immersed in cold NP-40 protein lysis buffer (Tris-HCL(pH7.6), 150mM NaCl, 1% Np-40) with Complete Protease Inhibitors Cocktail (Roche)) and sonicated using 5 pulses of 5 seconds at low output (2). Proteins lysate were centrifuged to remove non-dissolved proteins and followed by quantification using the Bradford protein assay (BioRad Laboratories). Immunoprecipitation protocol followed. Whole protein extracts from P13 and P15 mice (4 retinas per age, 1mg total protein per IP) were prepared and incubated with the cone transducin (Gɑt2) antibody bound to Dynabeads (Invitrogen). As control, 1mg of P13 whole protein extract was incubated with rabbit IgG bound to Dynabeads. Following precipitation and washings, immunoprecipitated Gpsm2 was detected on Western blots with Rabbit anti-GPSM2 antibody. For GPSM2/Gnat2 in vitro IP assay, 293T cells were co-transfected with vectors driving the expression of GFP and flag- fusion constructs. Total protein extract was obtained by applying NP40 lysis buffer on cells 24 hours post transfection. Proteins lysate were centrifuged to remove non-dissolved proteins and followed by quantification using the Bradford protein assay (BioRad Laboratories). Immunoprecipitation protocol followed. Whole protein extracts were incubated with the cone Flag antibody bound to Dynabeads (Invitrogen). Following precipitation and washings, immunoprecipitated GFP was detected on western blots with Rabbit anti-GFP antibody.

### Ultrastructure expansion microscopy of the mouse retina

Protocol for retina expansion was adapted from the Ultrastructural Expansion Microscopy (U-ExM) method^32^ with few optimizations. Crosslinking prevention step was extended to overnight (ON) incubation with 500 μL of 2% acrylamide (AA; A4058, Sigma-Aldrich) + 1.4% formaldehyde (FA; F8775, Sigma-Aldrich) at 37°C in an Eppendorf tube for each retina. The next day, the crosslinking solution was discarded and 200 μL of monomer solution (MS) composed of sodium acrylate (19% from 38% (w/w) stock solution diluted with nuclease-free water, 408220, Sigma-Aldrich), Acrylamide solution (10% from 40% stock solution A4058, Sigma-Aldrich), 0,1% N, N′- methylenbisacrylamide (BIS, from 2% stock, M1533, Sigma-Aldrich), 20μL of PBS 10X and completed to a total of 200 μL of nuclease-free water was added for 90 min at RT. MS was discarded and replaced with 350 μL of MS was added together with 0.02% ammonium persulfate (APS, 17874, Thermo Fisher Scientific) and 0.00006% tetramethylethylenediamine (TEMED, 17919, Thermo Fisher Scientific) for 45 min at 4°C first followed by 3 h incubation at 37°C to allow gelation. For this step, the retina was transferred into a small 0.27”x0.27 plastic mold (22-038-271, ThermoFisher) and covered with a coverslip coated with vaseline was added on top to close the chamber. Next, the coverslip was removed and 1 mL of denaturation buffer (200 mM SDS, 200 mM NaCl, 50 mM Tris Base in water (pH 9)) was added into the mold for 15 min at RT with shaking. Then, the gel was careful detached from the dish with a spatula and incubated in a 15 mL falcon tube filled with denaturation buffer for 1 h at 95°C and then ON at RT with shaking. Then, the gel was cut around the retina and expanded in 3 successive ddH2O baths. The gel was then manually sliced with a razorblade to obtain approximately 0.5 mm thick transversal sections of the retina that were then processed for immunostaining. PBS was discarded and replaced by 3 baths of PBS1X for 15 minutes each. After transfer into a 48-well plate, 500μL of blocking solution

(PBS/BSA3% + Thimerosal) was added to the gel for 2hrs with agitation at 37°C. The blocking solution was then replaced with the solution containing the primary antibodies diluted in blocking solution and incubated overnight at 37°C with agitation. Gels were washed 3 times 10 min at RT with PBS + Tween20 0.1%, (PBST) under agitation. After last wash, blocking solution containing secondary antibodies & Hoechst were added to the gels (500 μL / gel) and incubated overnight at 37°C under agitation. Gels were than washed 3 times using PBST and immersed in a large volume of distilled water for a last round of expansion overnight before acquisition with an inverted confocal microscope (LSM700, Zeiss) the next day. For pan-protein staining of expanded retinas, an additional step was added by immersion with Bodipy FL NHS Ester (Succinimidyl Ester; Thermo Scientific cat#D2184) in PBS 1X (10μM) at room temperature for 1h on gentle rotation. Mentioned scale bars were adjusted for an expansion factor of 4.2 based using rod nuclei diameter as fiducial.

### Statistical Analysis

Quantifications are reported as mean ± SD. Statistical tests are two-tailed unpaired t-test or one-way ANOVA with Tukey#s multiple comparison test or two-way ANOVA with post-hoc multiple comparison test as indicated in figure legends. t-test was adjusted with Bonferroni corrections, when necessary, as indicated. The p-values reported in each figure legend are labeled as follows: *p < 0.05, **p < 0.01, and ***p <

0.001. Reported *n* values represent the number of biological replicates (independent animals).

## Resource availability

### Lead contact

Further information and requests for resources and reagents should be directed to and will be fulfilled by the lead contact, Michel Cayouette

### Materials availability

Plasmids generated in this study are available upon request. Requests for plasmids should be directed to and will be fulfilled by the lead contact.

### Data and code availability

- Microscopy data reported in this paper will be shared by the lead contact upon request.
- This paper does not report original code.
- Any additional information required to reanalyze the data reported in this work paper is available from the lead contact upon request.

