## Supplemental information for "A non-canonical planar cell polarity pathway triggered by light"

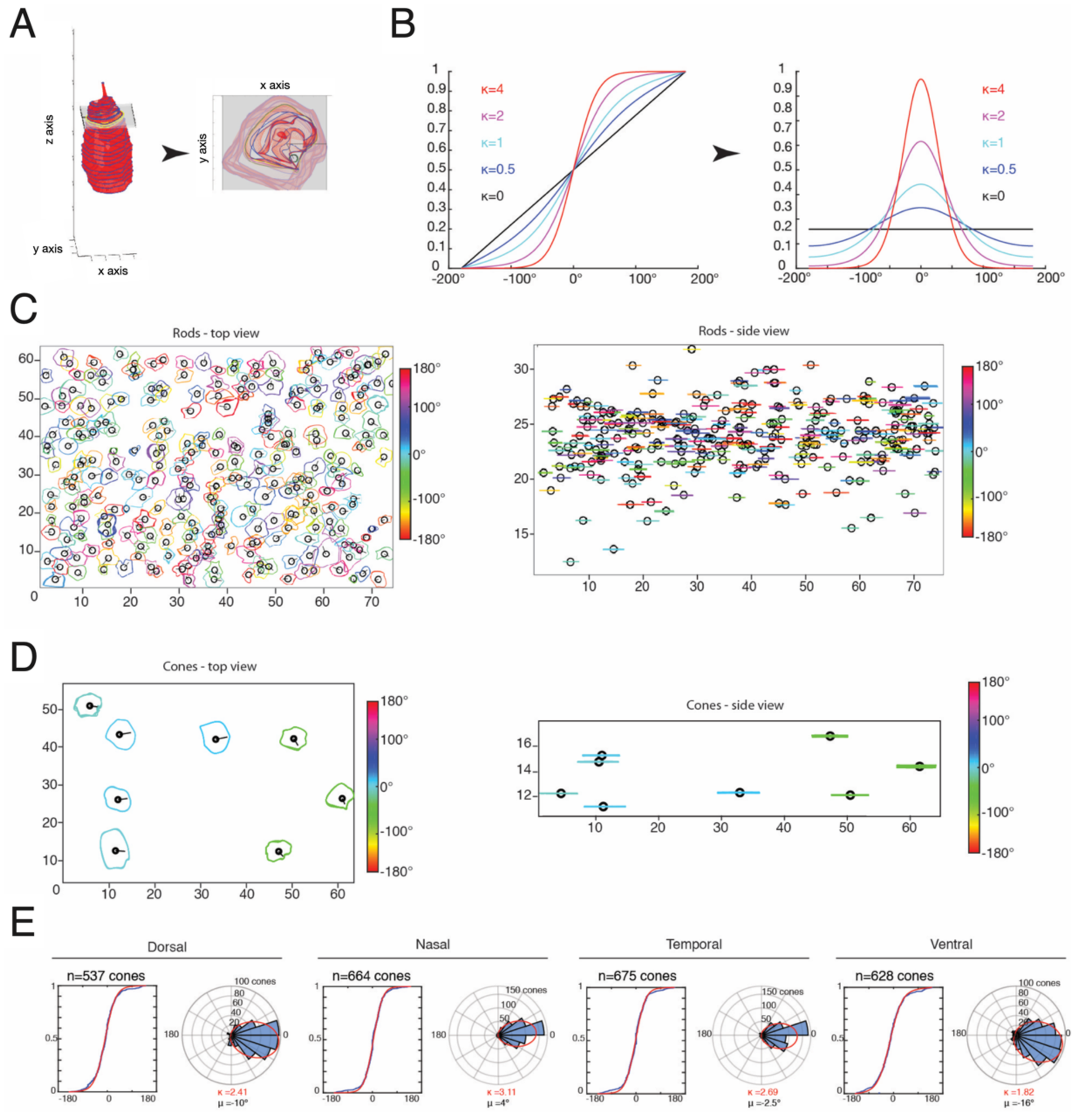


**Figure S1. FIBSEM segmentation and analysis of PCP in the adult mouse retina**
**(A)** Segmented cone IS reconstructed in 3D used for basal body analysis. The IS membrane border is interpolated (red lines) between the segmented IS (blue lines). Panel on the right shows a top view of the sliced plane shown on the left (grey) in the basal body region. The mean radius of the IS (yellow border) and the basal body (green) orientations are used for analysis. **(B)** Cumulative distribution (left) and probabilistic distribution (right) of Von Mises functions with different kappa values. **(C-D)** Top view (left) and side view (right) of rod (c) and cone (d) IS membranes (colored) and basal body (black circle) color-coded for the α value in en-face FIBSEM acquisition of wildtype retinas. **(E)** Cumulative distribution (left) and polar histogram (right) of all basal body orientations measured with the associated Von Mises function fit (red line) in different quadrants of wild-type retinas. Associated kappa (red) and mu (black) values are shown. The total number (*n*) of cones analyzed from 6 independent mice for each region is indicated on the top of each graph.

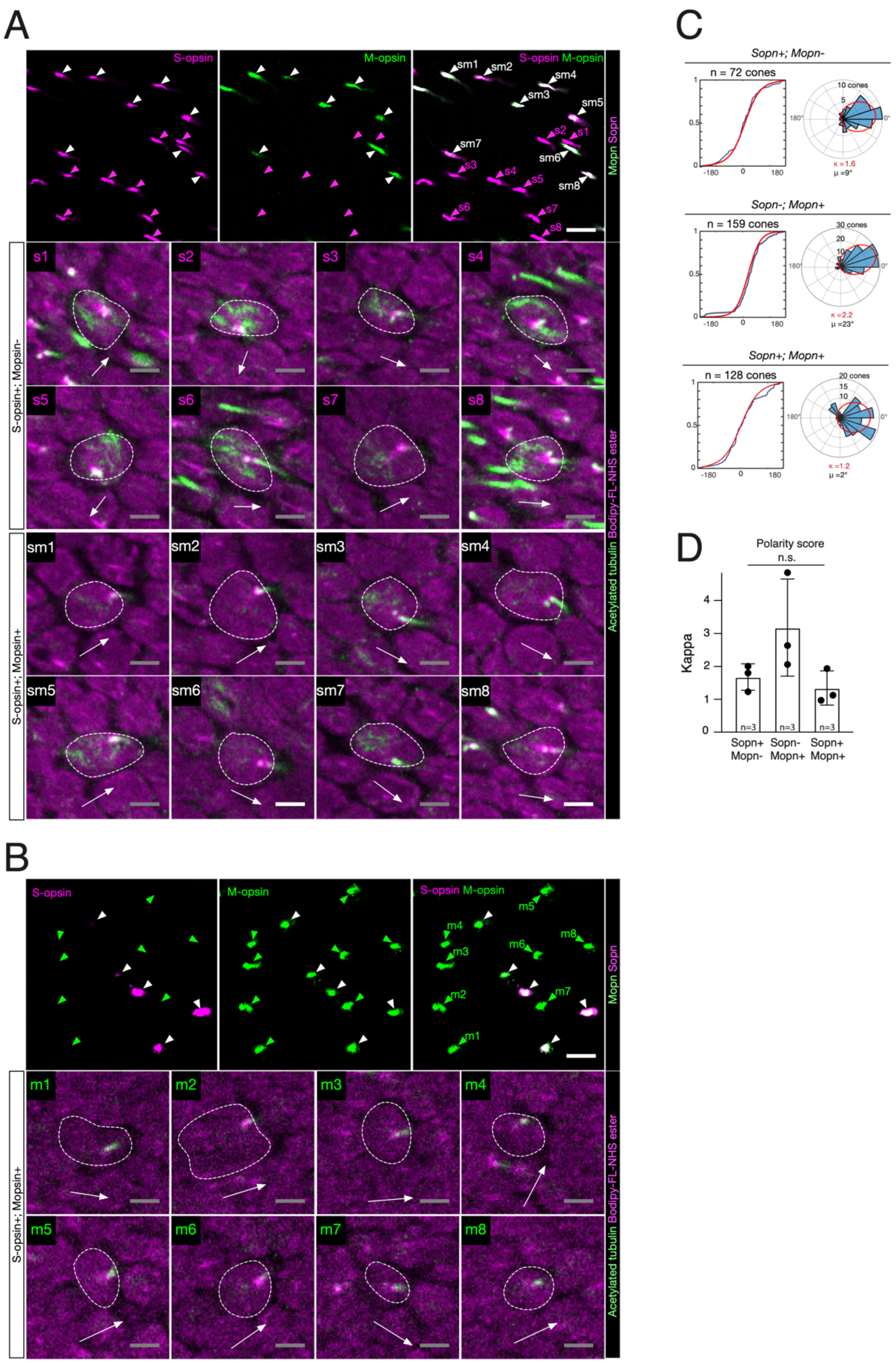


**Figure S2. Genuine cone exhibit PCP in the adult mouse retina**

(**A,B**) Upper panel : Confocal acquisitions in the outer segment plane of expanded adult mouse retina stained for Short-wavelength opsin (S-opsin, magenta) and Mediumwavelength opsin (M-opsin, green). Merged image (Right) indicates Sopn+;Mopn- cones (magenta arrowheads), Sopn-;Mopn+ cones (green arrowheads) and Sopn+;Mopn+ cones (white arrowheads). Lower panel : Confocal acquisitions in the basal body region of the corresponding cones identified in the upper panel stained for basal body and microtubules (acetylated tubulin, green) and a pan-protein marker outlining photoreceptors inner segments (Bodipy-FL-NHS, magenta). For each cone, orientation of the basal body is indicatd (white arrow). **(C)** Cumulative distribution (left) and polar histogram (right) of basal body planar orientation in cones and associated Von Mises distribution function fit (red line) of the corresponding cone cell type based on opsin expression. Associated kappa (red) and mu (black) values are shown. The total number (n) of cones analyzed from 3 independent mice is indicated on top of each graph. **(D)** Histogram of cone polarity scores in *the different cone types*. Total number (*n*) of cones analyzed: *Sopn+;Mopn-* = 72, *Sopn-;Mopn+* = 159, *Sopn+;Mopn+ = 128 cones*. One-way ANOVA: p=0.11. Scale bars: 10μm (A upper panel), 2μm (lower panel zoomed-in insets.

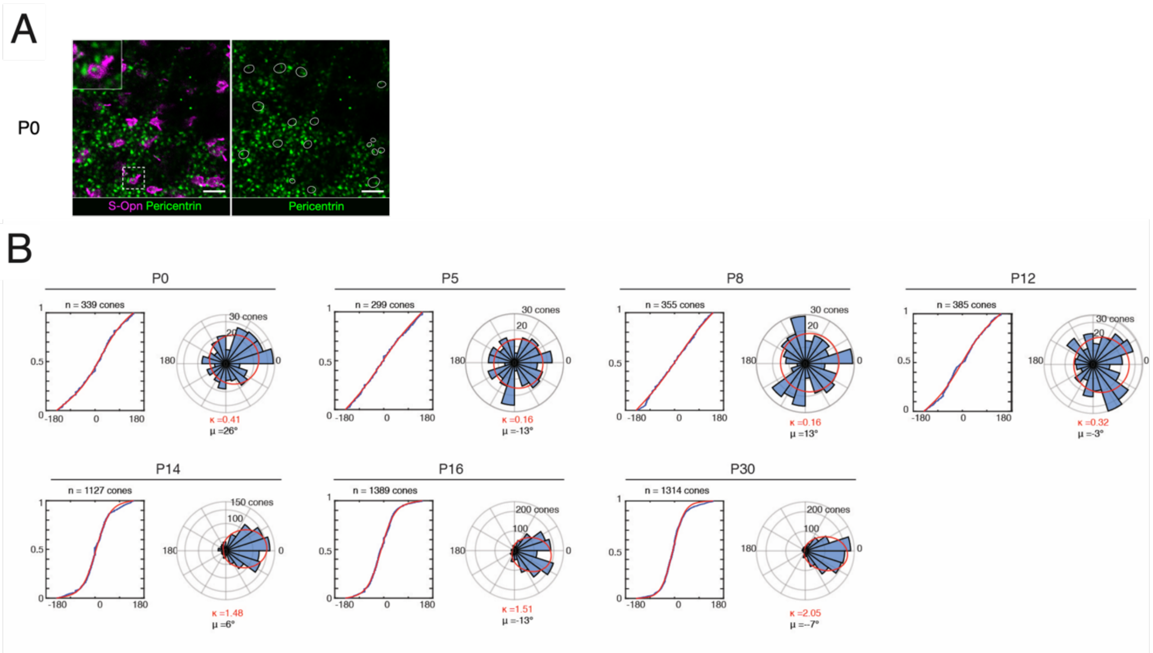


**Figure S3. Cone PCP analysis during development**

(**A**) Confocal acquisition of flat mount of P0 retina stained for pericentrin (green), S-Opsin (Magenta). Cones are outlined by a white line. At P0, basal bodies are centrally located (zoomed-in insets). **(B)** Cumulative distribution (left) and polar histogram (right) of cone basal body orientations and the associated Von Mises function fit (red line) at different ages. Associated kappa (red) and mu (black) values are shown. The total number (*n*) of cones analyzed from 3 mice (P14, P16, P30); 4 mice (P0, P5, P8); and 5 mice (P12) is indicated on top of each graph. P: post-natal stage in days.

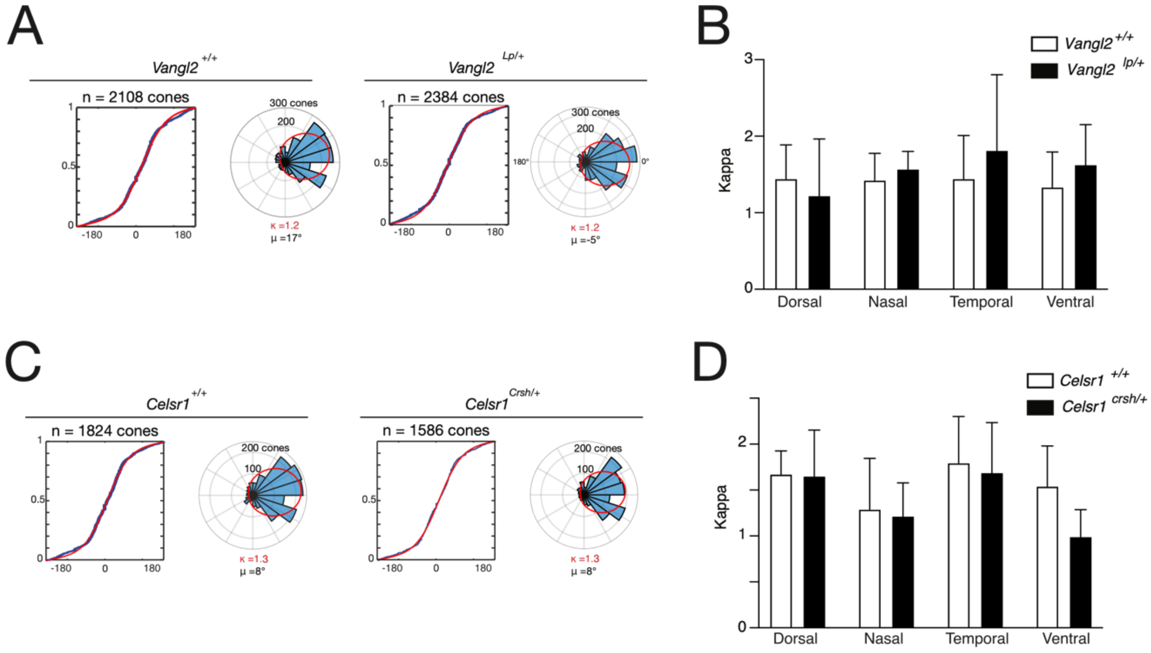


**Figure S4. Cone PCP analysis in retina of mice mutants of the core PCP pathway**
**(A)** Cumulative distribution (left) and polar histogram (right) of basal body orientations and associated Von Mises distribution function fit (red line) of *Vangl2*$^{+/+}$ and *Vangl2*$^{Lp/+}$ mice. Associated kappa (red) and mu (black) values are shown. The total number (*n*) of cones analyzed from 5 independent mice is indicated on top of each graph. **(B)** Histogram of cone polarity scores in *Vangl2*$^{+/+}$ and *Vangl2*$^{Lp/+}$ mice for each different region. Two-way ANOVA: Petal factor p=0.60, Genotype factor: p=0.59. **(C)** Cumulative distribution (left) and polar histogram (right) of basal body orientations and associated Von Mises distribution function fit (red line) of *Vangl2*$^{+/+}$ and *Vangl2*$^{Lp/+}$ mice. Associated kappa (red) and mu (black) values are shown. The total number (*n*) of cones analyzed from 4 independent mice is indicated on top of each graph. **(D)** Histogram of cone polarity scores in *Celsr1*$^{+/+}$ and *Celsr1*$^{Crsh/+}$ mice for each different region. Two-way ANOVA: Petal factor p=0.13, Genotype factor: p=0.06.

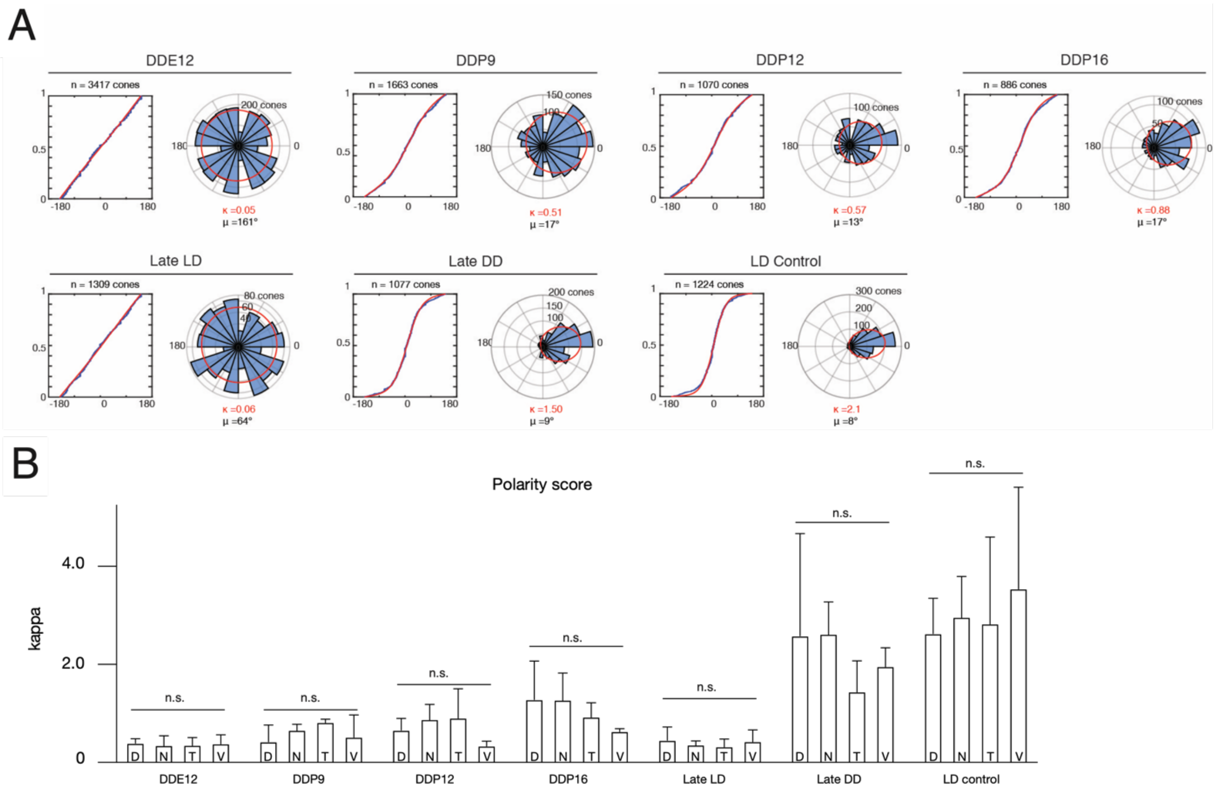


**Figure S5. Cone PCP analysis after different dark-rearing protocols**

(**A**) Cumulative distribution (left) and polar histogram (right) of basal body orientations and the associated Von Mises function fit (red line) in the different dark-reared experiments. Associated kappa (red) and mu (black) values are shown. The total number (*n*) of cones analyzed from at least 3 independent mice for each region is indicated on top of each graph. Mice reared in constant darkness (dark:dark; DD) from embryonic day (E) 12.5 (DDE12), P9 (DDP9), P12 (DDP12) or P16 (DDP16) onward and sacrificed at P30 for analysis. Late light:dark (LD) : mice reared in constant darkness between E12.5 and P22 and switched to 12 hours light-dark cycles at P23 onward until sacrifice at P360. Late dark:dark (DD) : mice reared in constant darkness between P120 and sacrifice at P150. (**B**) Histogram of cone polarity scores in the different quadrants of the retina from mouse reared in different lightning conditions. Two-way ANOVA: Petal factor p=0.8, Dark-rearing condition factor: p<0.0001, post-hoc mutlicomparison mixed-effects comparing petals not significant (n.s.).

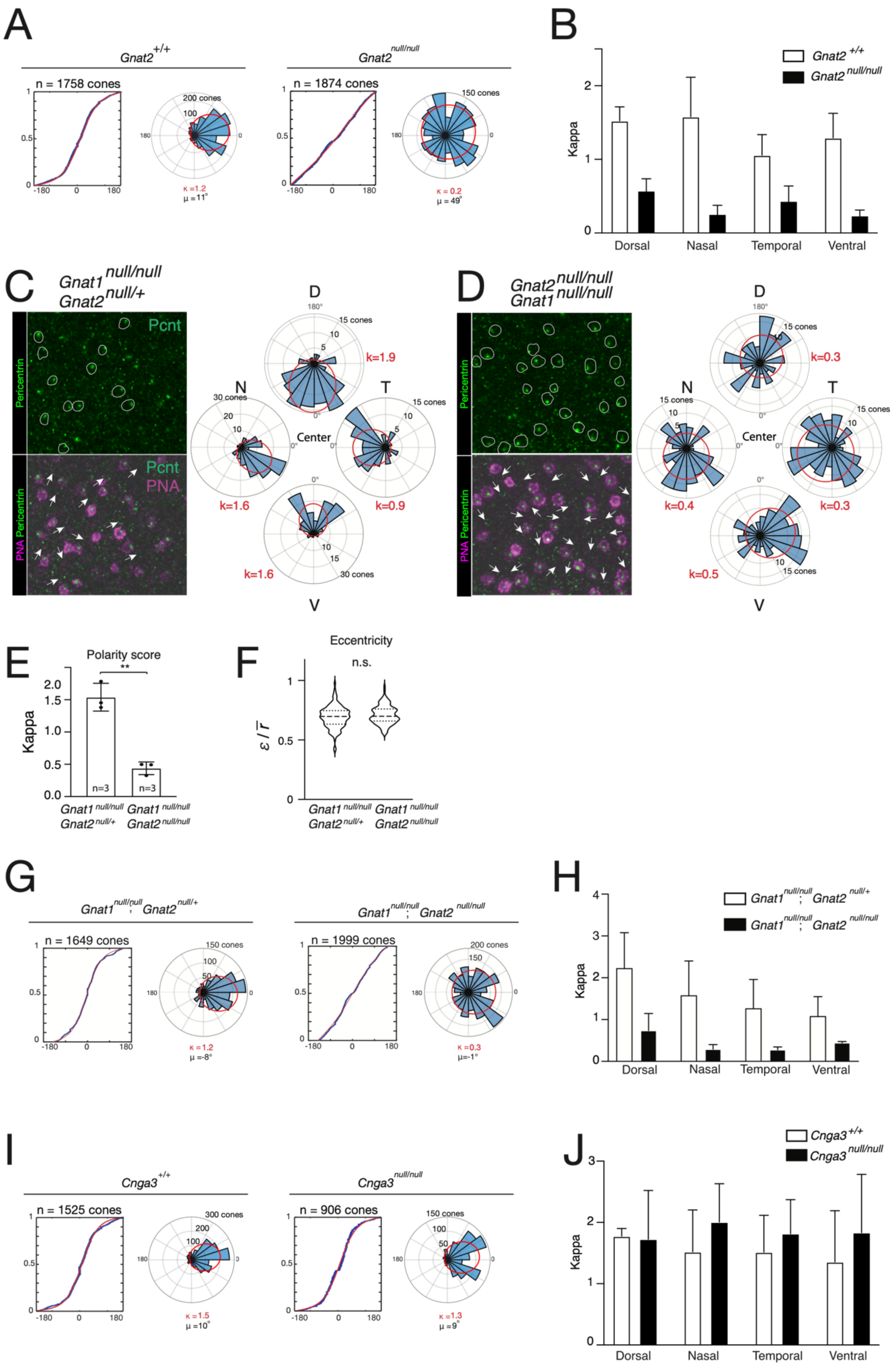
A
Gnat2 +/+
n = 1758 cones
κ =1.2
μ =11°
Gnat2 null/null
n = 1874 cones
κ = 0.2
μ = 49°
B
Gnat2 +/+
Gnat2 null/null
Kappa
Dorsal
Nasal
Temporal
Ventral
C
Gnat1 null/null
Gnat2 null/+
Pericentrin
PNA Pericentrin
Pcnt
PNA
D
N
T
V
Center
k=1.9
k=1.6
k=0.9
k=1.6
D
Gnat2 null/null
Gnat1 null/null
k=0.3
k=0.4
k=0.3
k=0.5
E
Polarity score
**
Kappa
n=3
Gnat1 null/null Gnat2 null/+
Gnat1 null/null Gnat2 null/null
F
Eccentricity
n.s.
ε / r̄
G
Gnat1 null/null; Gnat2 null/+
n = 1649 cones
κ =1.2
μ =-8°
Gnat1 null/null; Gnat2 null/null
n = 1999 cones
κ =0.3
μ=-1°
H
Gnat1 null/null; Gnat2 null/+
Gnat1 null/null; Gnat2 null/null
Kappa
Dorsal
Nasal
Temporal
Ventral
I
Cnga3 +/+
n = 1525 cones
κ =1.5
μ =10°
Cnga3 null/null
n = 906 cones
κ =1.3
μ =9°
J
Cnga3 +/+
Cnga3 null/null
Kappa
Dorsal
Nasal
Temporal
Ventral

**Figure S6. Cone PCP analysis in mouse mutants of genes coding for effectors of the phototransduction cascade**

**(A)** Cumulative distribution (left) and polar histogram (right) of basal body orientations and associated Von Mises function fit (red line) of *Gnat2*$^{+/+}$ and *Gnat2*$^{null/null}$ mice. Associated kappa (red) and mu (black) values are shown. The total number (*n*) of cones analyzed from 3 independent mice is indicated on top of each graph. **(B)** Histogram of cone polarity scores in *Gnat2*$^{+/+}$ and *Gnat2*$^{null/null}$ mice. Two-way ANOVA: Petal factor p=0.12, Genotype factor: p<0.0001. **(C-D)** Confocal acquisitions of *Gnat1*$^{null/null}$; *Gnat2*$^{null/+}$ (C) and *Gnat1*$^{null/null}$; *Gnat2*$^{null/null}$ (D) retina flat-mounts stained for pericentrin (Pcnt, green) and peanut agglutinin (PNA, magenta). Representative examples of cones (outlined) with various centrosome orientation (arrows) are shown. For each genotype, a representative rose plot of basal body orientation (α-angles distribution) and the associated kappa value is shown. **(E)** Histogram of cone polarity scores in *Gnat1*$^{null/null}$; *Gnat2*$^{null/+}$ and *Gnat1*$^{null/null}$; *Gnat2*$^{null/null}$ mice. Total number (*n*) of cones analyzed: *Gnat1*$^{null/null}$; *Gnat2*$^{null/+}$ = 1649, *Gnat1*$^{null/null}$; *Gnat2*$^{null/null}$ = 1999 cones. Two-tailed unpaired t-test: p=0.0013 (**p<0.01). **(F)** Violin plots of cone basal body eccentricities for each genotype (n=3 animals, 360 cells counted for each genotype). Two-tailed nested unpaired t-test: p=0.09 **(G)** Cumulative distribution (left) and polar histogram (right) of basal body orientations and associated Von Mises function fit (red line) of *Gnat1*$^{null/null}$; *Gnat2*$^{null/+}$ and *Gnat1*$^{null/null}$; *Gnat2*$^{null/null}$ mice. Associated kappa (red) and mu (black) values are shown. The total number (*n*) of cones analyzed from 3 independent mice is indicated on top of each graph. **(H)** Histogram of cone polarity scores in *Gnat1*$^{null/null}$; *Gnat2*$^{null/+}$ and *Gnat1*$^{null/null}$; *Gnat2*$^{null/null}$ mice in the different quadrants of the mouse retina. Two-way ANOVA: Petal factor p=0.10, Genotype factor: p=0.0001. **(I)** Cumulative distribution (left) and polar histogram (right) of basal body orientations and associated Von Mises function fit (red line) of *Cnga3*$^{+/+}$ and *Cnga3*$^{null/null}$ mice. Associated kappa (red) and mu (black) values are shown. The total number (*n*) of cones analyzed from 3 independent mice is indicated. **(J)** Histogram of cone polarity scores in *Cnga3*$^{+/+}$ and *Cnga3*$^{null/null}$ mice in the different quadrants of the mouse retina. Two-way ANOVA: Petal factor p=0.97, Genotype factor: p=0.27.

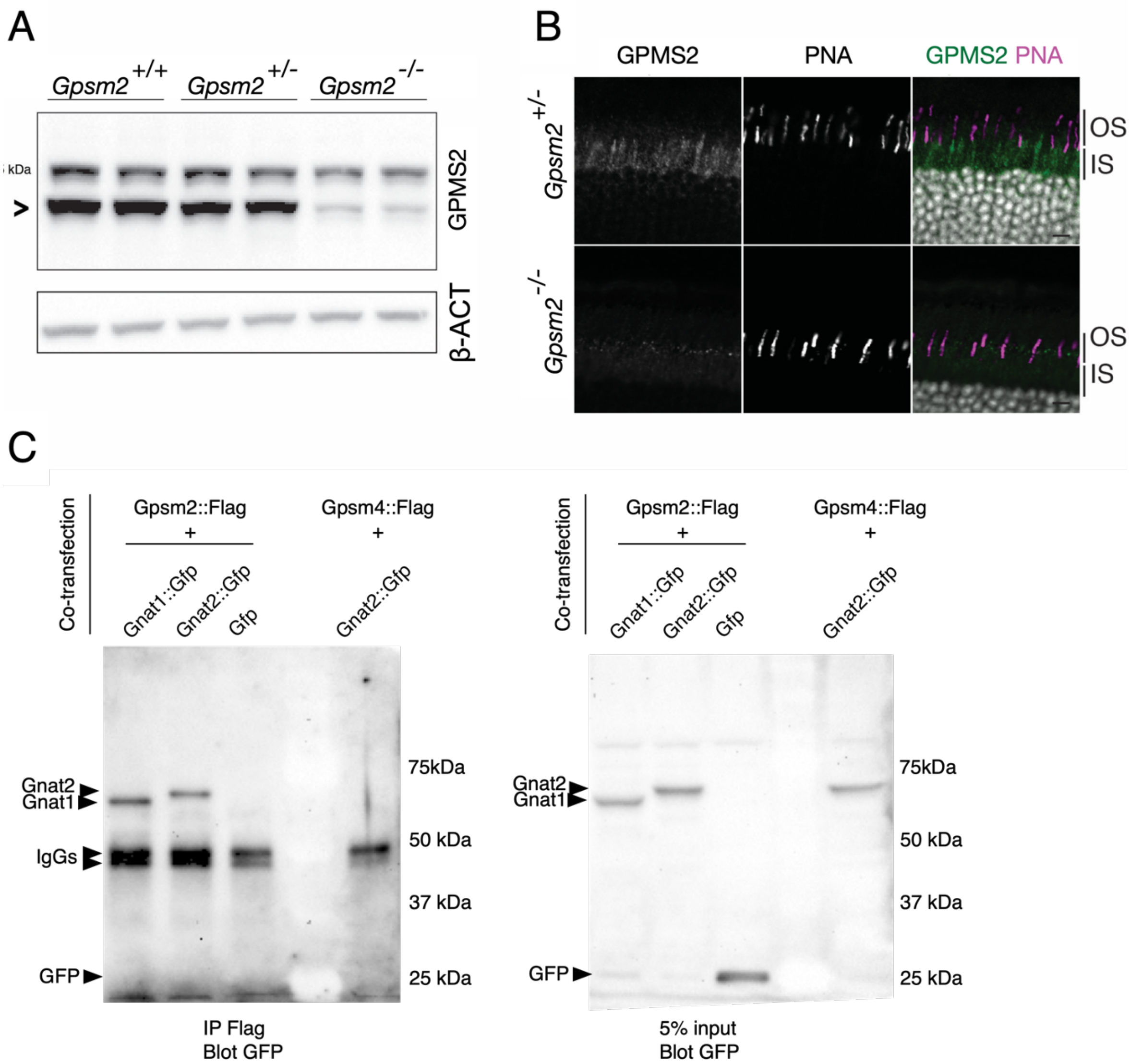


**Figure S7. Anti-GPSM2 antibody and GPSM2-G$\alpha$t2 *in vitro* interaction validations**
**(A)** Western blot on total adult retina extracts from *Gpsm2*$^{+/+}$, *Gpsm2*$^{+/-}$, and *Gpsm2*$^{-/-}$ mice using the rabbit anti-GPSM2 antibody (Sigma). Observing the specific GPSM2 band (arrowhead) we noted a decrease in intensity when comparing protein extracts from *Gpsm2*$^{+/+}$, *Gpsm2*$^{+/-}$ and *Gpsm2*$^{-/-}$ mice. Notably, in the *Gpsm2* KO extract, a faint yet discernible band persisted, suggesting a weak affinity of the antibody for a non-specific protein of similar size to GPSM2. **(B)** Immunohistochemistry for GPSM2 (goat antibody from Life Technology) and peanut agglutinin (PNA) on adult retinal sections from *Gpsm2*$^{+/-}$ and *Gpsm2*$^{-/-}$. Note that the GPSM2 signal in the inner segments of photoreceptors is almost totally lost in the knockout mice, with a faint remaining signal, validating the specificity of this antibody for staining. (**C**) Green fluorescent Protein (GFP) blot after Flag immunoprecipitation (left) or 5% input (right) from protein extracts of 293 cells co-transfected with the indicated expression vector plasmids. While Gpsm2:Flag could co-immunoprecipitate Gnat1:GFP (positive control) and Gnat2:GFP (experimental), it could not co-IP GFP. Conversely, GPSM4:Flag could not IP Gnat2:GFP, demonstrating specificity of the GPSM2-Gnat2 pull-down. Scale bar:10µm.

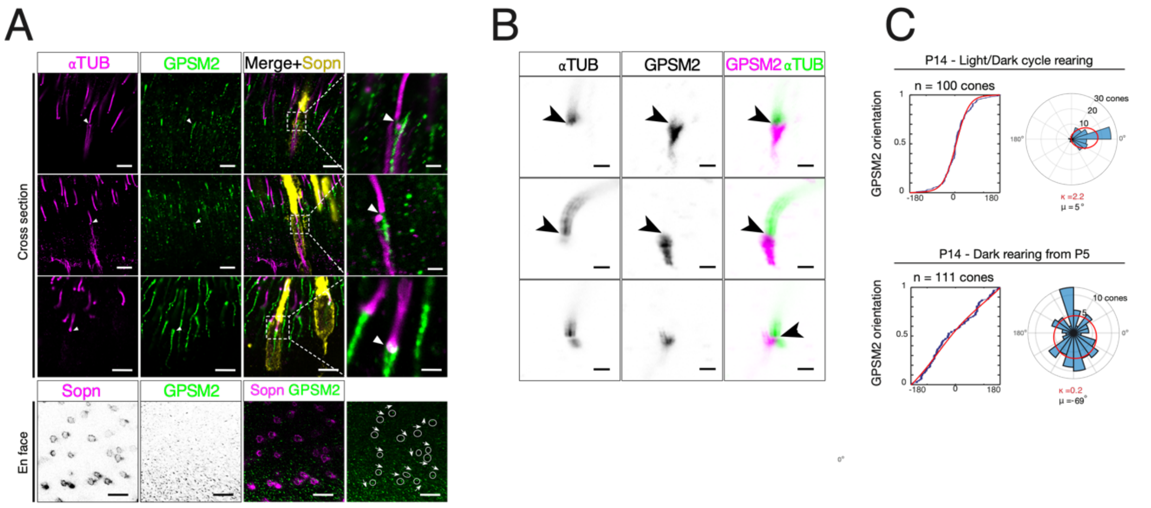


**Figure S8. Planar polarization of GPSM2 immunostaining signal in mouse cones**
**(A)** Optical section acquisitions of mouse retina after ultrastructure expansion and immunostaining for acetylated tubulin (αTUB, magenta, upper panel), GPSM2 (green) and S-Opsin (yellow, upper panel; magenta, lower panel). Upper panel: Arrowhead point to the distal GPSM2 in the basal body region of cones. Lower panel: representative example of cones (outlined) with GPSM2 planar orientations (arrows) are shown. **(B)** Confocal acquisitions of P13 mouse photoreceptors after ultrastructure expansion immuno-labeled for acetylated tubulin (αTUB, green) and GPSM2 (magenta). GPSM2 form processes that come in contact with the base of the distal region of the basal body (arrowhead). **(C)** Cumulative distribution (left) and polar histogram (right) of GPSM2 planar orientation in cones and associated Von Mises function fit (red line) of P14 expanded retina from mice reared in light/dark cycles (upper) or constant darkness from P5 (lower). Associated kappa (red) and mu (black) values are shown. The total number (n) of cones analyzed from 3 independent mice is indicated on top of each graph. Scale bars: 5µm (A upper panel), 1µm (A zoomed-in insets), 10µm (A lower panel), 500nm (B).

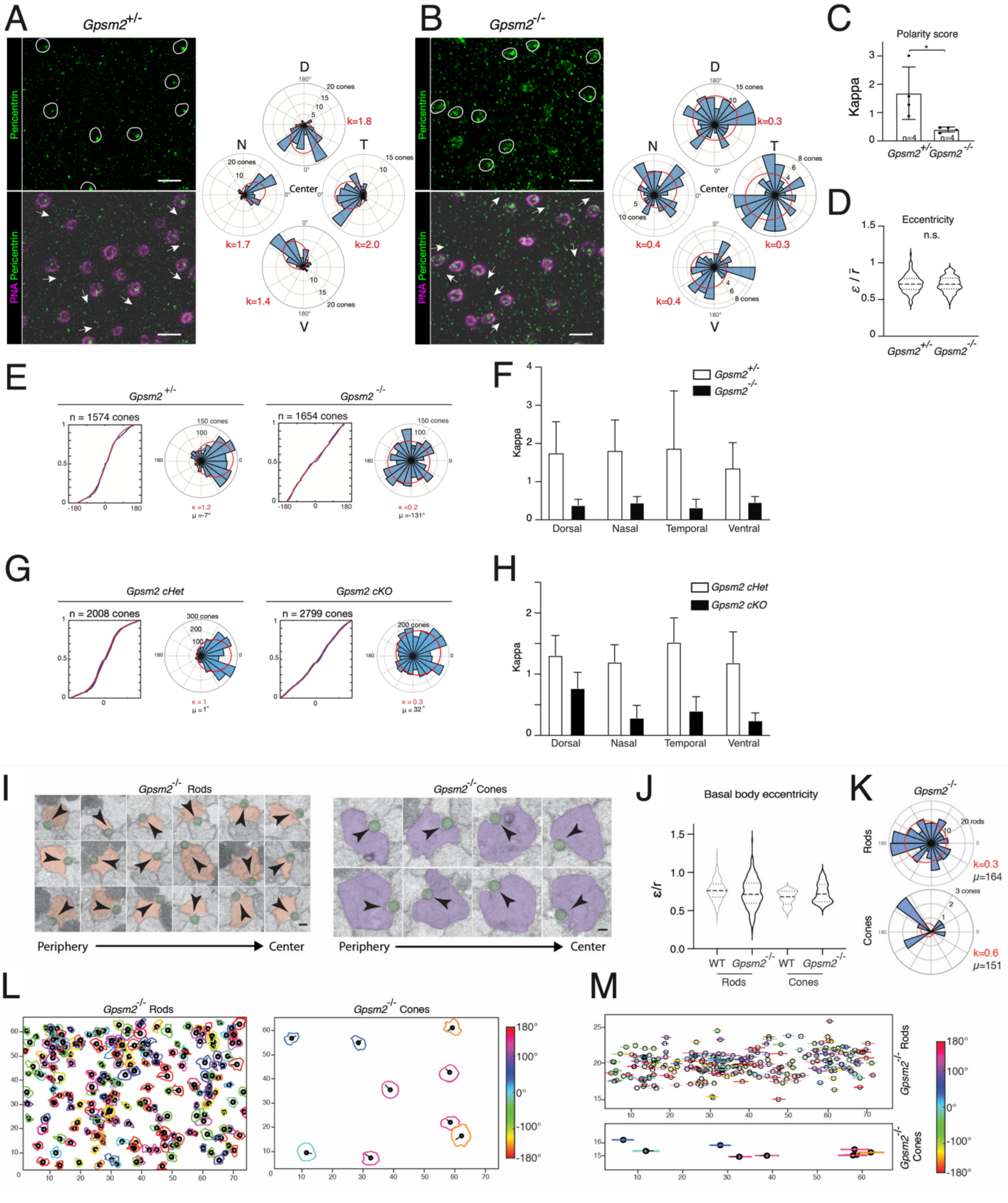


**Figure S9. Cone PCP analysis in the Gpsm2 mutant mice**

**(A, B)** Confocal acquisitions of *Gpsm2*$^{+/-}$ (A) and *Gpsm2*$^{-/-}$ (B) retina flat mounts stained for pericentrin (PCNT, green) and Peanut agglutinin (PNA, magenta). Representative examples of cones (outlined) with various basal body orientations (arrows) are shown. For each genotype, a representative rose plot of basal body orientations (α-angles distribution) and the associated kappa value is shown. Dorsal (D); Nasal (N); Temporal (T); Ventral (V). scale bars: 10µm. **(C)** Histogram of cone polarity scores in *Gpsm2*$^{+/-}$ and *Gpsm2*$^{-/-}$ mice. Graph shows mean ±s.d. *n* indicates biological replicates. Total number of cones analyzed: *Gpsm2*$^{+/-}$ = 1574 cones, *Gpsm2*$^{-/-}$ = 1654 cones. Two-tailed unpaired t-test p=0.03 (*p<0.05). **(D)** Violin plots of basal body eccentricities (n=3 animals, 360 cells counted for each genotype). Two-tailed unpaired t-test: p=0.62. **(E)** Cumulative distribution (left) and polar histogram (right) of basal body orientations and associated Von Mises function fit (red line) of *Gpsm2*$^{+/+}$ and *Gpsm2*$^{null/null}$ mice. Associated kappa (red) and mu (black) values are shown. The total number (*n*) of cones analyzed from 4 independent mice is indicated on top of each graph. **(F)** Histogram of cone polarity scores in *Gpsm2*$^{+/+}$ and *Gpsm2*$^{null/null}$ mice in the different quadrants of the mouse retina. Two-way ANOVA: Petal factor p=0.93, Genotype factor: p<0.0001. **(G)** Cumulative distribution (left) and polar histogram (right) of basal body orientations and associated Von Mises function fit (red line) of *Gpsm2* cHet and *Gpsm2* cKO mice. Associated kappa (red) and mu (black) values are shown. The total number (*n*) of cones analyzed from 4 independent mice is indicated on top of each graph. **(H)** Histogram of cone polarity scores in *Gpsm2* cHet and *Gpsm2* cKO mice in the different quadrants of the mouse retina. Two-way ANOVA: Petal factor p=0.07, Genotype factor: p<0.0001. **(I)** High magnification FIBSEM images of randomly selected rods left (red), and cones (right, purple) in the connecting cilium region of dorsal retina region from a P40 *Gpsm2*$^{-/-}$ mice are shown. Orientation of the cilium (green) is indicated relative to the center of rod and cone IS by an arrowhead. Scale bar: 500nm. **(J)** Violin plots of rod and cone basal body eccentricity measured in FIBSEM acquisition of wild-type (WT, grey) and *Gpsm2*$^{-/-}$ mice (black). Wild-type: n=280 rods; n=8 cones; *Gpsm2*$^{-/-}$: n=241 rods; n=9 cones. One-way ANOVA p=0.07. **(K)** Polar histograms of basal body planar orientations (blue bins) and the associated density probability fit (red line). Associated kappa (red) and mu (black) values are shown. **(L, M)** Top view (L) and side view (M) of rod and cone IS membranes (colored) and basal body (black circle) color-coded for the α angle value for the FIBSEM acquisition of *Gpsm2*$^{-/-}$ mice. Scale bar: 500nm.

**Supplementary Movie S1. Mouse photoreceptors segments tridimensional ultrastructure**

Slice-and-view from different angles of FIBSEM images of the wild-type mouse retina acquired in cross-section orientation. Segmentation and reconstruction of rod IS (light green) and cones IS (magenta) show localization of the connective cilium (red).
